# Protein Malnutrition Facilitates Intestinal Colonization with Highly Resistant *Klebsiella pneumoniae*

**DOI:** 10.1101/2025.07.21.665917

**Authors:** Thomas Holowka, Veronica Feijoli Santiago, Jamie Xiao, Mera F. Liccione, Alisher Bimagambetov, Kenneth Walsh, Jason W. Arnold, Kalisvar Marimuthu, Oon Tek Ng, Jonathan R. Swann, David van Duin, Luther A. Bartelt

## Abstract

Pediatric infections with Highly Resistant Enterobacterales (HRE), including *Klebsiella pneumoniae* resistant to 3^rd^-generation cephalosporins and/or carbapenems, disproportionately affect low- and middle-income countries where malnutrition is prevalent. The underlying mechanisms linking malnutrition to HRE colonization in children have not been established. In this study we developed a mouse model of pediatric malnutrition and intestinal colonization with clinical isolates of carbapenem-resistant *K. pneumoniae* (CR-Kp). Juvenile mice fed a protein-deficient diet (PD) were more susceptible to intestinal colonization after inoculation with human-derived strains of CR-Kp, demonstrating a 3-4 log higher colonization burden in comparison to mice fed a control diet (CD). Colonization in PD-fed mice persisted for up to 6 weeks and CR-Kp were transmitted between PD-fed but not CD-fed cage mates. Antibiotic treatment resulted in similar CR-Kp colonization burdens regardless of diet, suggesting that nutrition-dependent colonization resistance is reliant on an intact microbiota. Secondary bile acids, a product of resident intestinal microbiota, were reduced in PD-fed and antibiotic treated mice and demonstrated an inverse correlation with CR-Kp burden. Secondary bile acids directly inhibited CR-Kp growth *in vitro*, suggesting that a loss of these inhibitory metabolites may mediate malnutrition-induced susceptibility to HRE colonization.

## Introduction

Antimicrobial resistant bacteria present an urgent global threat responsible for over 1 million deaths worldwide in 2019, with mortality predicted to increase to nearly 2 million deaths annually by 2050 (1, 2). Among the greatest contributors to the global toll of antimicrobial resistance are highly-resistant Enterobacterales (HRE) including species of *Klebsiella pneumoniae* and *E. coli* resistant to 3^rd^ generation cephalosporins and carbapenems (2). HRE colonize the intestine where they can persist silently for months to years prior to progressing to invasive infection in certain susceptible individuals (3–7). Healthcare exposure and antimicrobial use have been linked to HRE colonization (8–10). However, individual host comorbidities that confer risk for colonization and infection remain incompletely understood. The highest rates of mortality attributed to HRE worldwide occur in low- and middle-income countries where malnutrition is also most prevalent (11–13). Malnutrition is a known risk factor for bloodstream bacterial infection in children residing in Sub-Saharan Africa and has been independently linked to HRE infection in this setting (14–16). Global guidelines recommend empiric antimicrobial administration in children with severe acute malnutrition, however near universal HRE colonization and increased transmission to household contacts has been reported in individuals receiving this treatment (17–19). Since malnutrition aflicts over 600 million individuals worldwide, and prolonged HRE carriage is linked with inter-individual spread, it is crucial to determine whether malnutrition facilitates HRE colonization independent from antimicrobial use and other healthcare exposures (12, 20). Furthermore, a mechanistic understanding of how malnutrition might impact colonization dynamics is crucial to curbing the global health impact of these high priority pathogens.

To establish colonization, HRE strains must access a niche within the complex community of microbes in the intestine (21, 22). The term “colonization resistance” refers to the propensity for the resident host microbiota to interfere with intestinal colonization by exogenous pathogens such as HRE (23, 24). Colonization resistance may be mediated through a number of activities including space and nutrient exclusion, antimicrobial metabolite generation, and stimulation of host defenses (25). Colonization resistance to intestinal HRE is most clearly illustrated by the impact of antimicrobials. Antimicrobial use is strongly linked to HRE colonization risk in hospitalized patients, and murine models demonstrate that antimicrobial-mediated depletion of the host microbiota facilitates intestinal colonization with HRE isolates (26–28). Nutritional protein deficiency can also profoundly re-shape the host microbiota leading to increased susceptibility to conventional diarrheal pathogens, such as diarrheagenic *E. coli* and parasites (29–32). Unlike antimicrobials, protein deprivation causes these changes without large-scale depletion of bacterial density in the intestine but rather through alterations in microbiome composition and metabolic output (33–35).

To determine if malnutrition directly facilitates HRE intestinal colonization acquisition, sustained carriage, and transmission independent of other exposures, we developed a new model of HRE colonization using several patient-derived isolates of carbapenem resistant *Klebsiella pneumoniae* (CR-Kp). In this model, juvenile mice fed a protein malnutrition diet demonstrated greater intestinal colonization and transmission of CR-Kp in a manner that was dependent on the CR-Kp isolate and correlated with loss of microbiota-generated inhibitory molecules in the intestinal microenvironment.

## Results

### CR-Kp colonization establishment and burden is dependent on intestinal microbiota disruption

Kp175, a representative of the ST307 OXA-48-producing lineage of CR-Kp that has global spread, was originally isolated from a patient with persistent rectal carriage for >100 days and documented transmission to household contacts (Table S1) (7). Inoculation of 10^6^ CFU of Kp175 via orogastric lavage into adult C57BL6 mice lead to colonization detected via shedding of 10-100 CFU/mg in feces (Fig 1A). Inconsistent or no colonization was observed in mice inoculated with 10^2^ or 10^4^ CFU (Fig 1B). Treatment of mice with continuous ampicillin in their drinking water reduced microbiota density ∼100 fold (Fig S1A). In ampicillin-treated mice there was consistent fecal shedding at a maximum density ∼10^7^ CFU/mg with an inoculating dose as low as 10^2^ CFU (Fig 1A, 1B, Table 1). Two other clinical strains of CR-Kp, ARLG-4605 and ARLG-4404 (Table S1) similarly achieved maximal colonization burden of ∼10^7^ CFU/mg in feces of mice on ampicillin, but were incapable of colonizing mice without ampicillin treatment (Fig 1C). After colonization establishment in mice on ampicillin, withdrawal of the antibiotic lead to a significant, but overall modest, 10-fold decline in colonization burden over a week after the antibiotic was withdrawn (Fig 1D). These findings demonstrate that adult mice with a fully intact microbiota resist CR-Kp, permitting colonization at a relatively low density only when administered at a high inoculating dose with select strains. In contrast, antibiotic-mediated depletion of the microbiota allows for a colonization at a much greater density that is largely fixed regardless of CR-Kp strain and inoculating dose, and that is persistent with only a small reduction even after antibiotic withdrawal.

**Figure 1:**
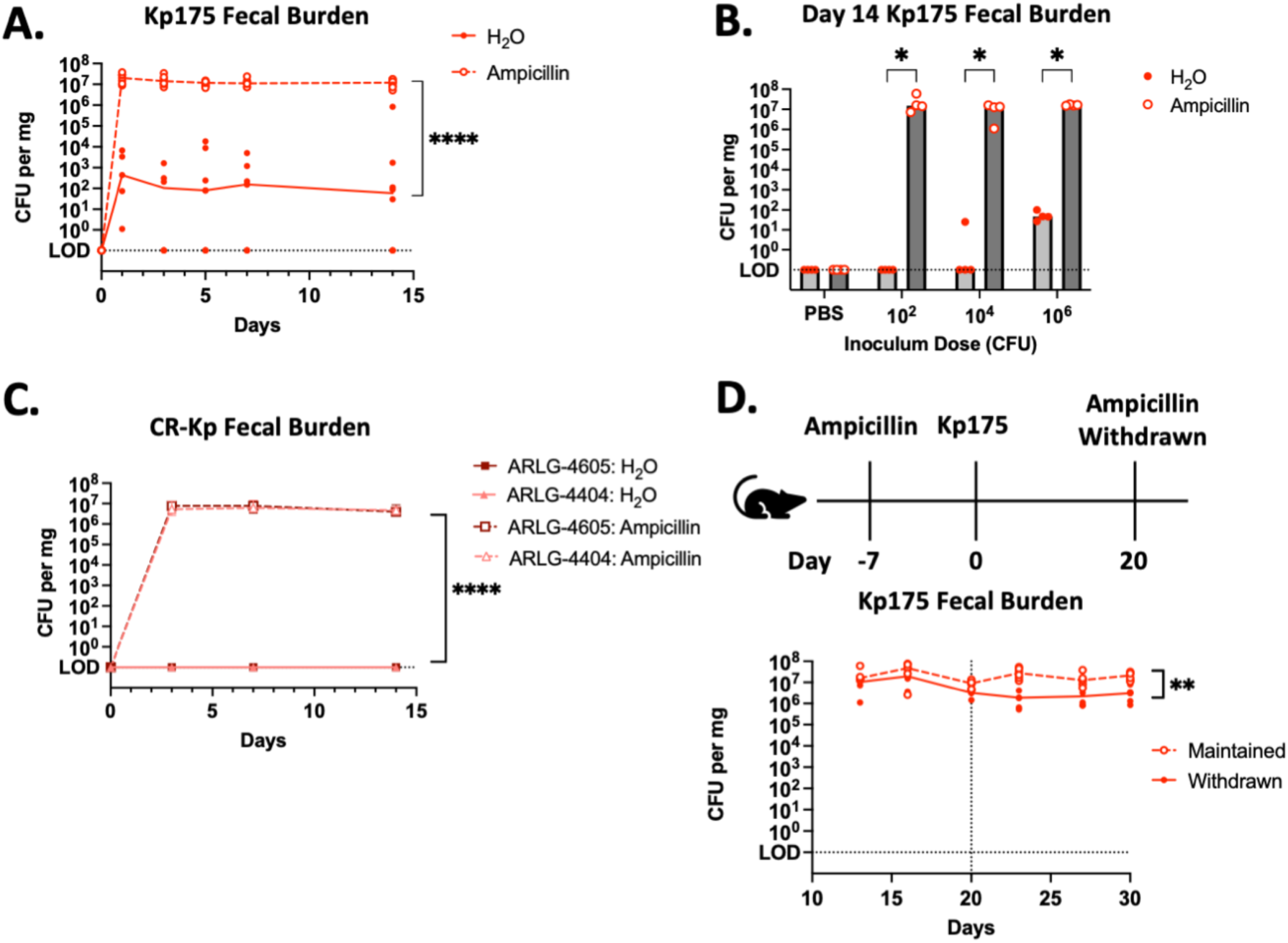
CR-Kp Colonization in Mice with or without Ampicillin Treatment. (A.) Mice were provided with regular drinking water (H_2_O) or with drinking water containing ampicillin 7 days prior to oral gavage with 10^6^ CFU Kp175. Fecal specimens were collected for culture-based quantification at days 0, 1, 3, 5, 14 after CR-Kp challenge. Median burden depicted, N=6 per group. 2-way ANOVA, ****=p<0.0001 (B.) Mice with or without ampicillin in their drinking water were challenged with PBS or Kp175 at the listed inoculum dose. Fecal specimens were collected for culture-based quantification at day 14. Median burden depicted, N=4 per group, Mann-Whitney U-test with 10% FDR, *= p<0.05. (C.) Mice with or without ampicillin in their drinking water were challenged with 10^6^ CFU ARLG-4605 or ARLG-4404. Fecal specimens were collected for culture-based quantification at days 0, 3, 7, 14 after CR-Kp challenge. Median burden depicted, N=6 per group. 2-way ANOVA, ****=p<0.0001 for ampicillin-treated versus regular drinking water for both ARLG-4605 and ARLG-4404 challenged mice. (D.) Mice with ampicillin in their drinking water were challenged with 10^6^ CFU Kp175, and in half of these mice ampicillin was replaced with regular water at day 20 after CR-Kp challenge. Fecal specimens were collected for culture-based quantification at days 12, 16, 20, 23, 27, 30 after CR-Kp challenge. Median burden depicted, N=4 per group. 2-way ANOVA, **=p<0.01.

**Table 1:** Kp175 Colonization Incidence and Peak Burden. Incidence of Kp175 colonization expressed as colonized/total (% colonized) and peak burden expressed as median CFU/mg in fecal specimens. Peak colonization burden (between day 7 and 14) reported in mice treated with or without ampicillin and in mice initiated on CD versus PD 7 days prior to inoculation with Kp175 at indicated inoculation dose across multiple experiments (A, B, C). P values shown for comparison of colonization incidence of PD-vs. CD-fed mice by Fisher’s Exact Test.

|  |  | 1.00E+02 |  | 1.00E+04 |  | 1.00E+06 |  | 1.00E+08 |  |
| --- | --- | --- | --- | --- | --- | --- | --- | --- | --- |
| H<sub>2</sub>O vs. Amp Treated |  | H<sub>2</sub>O | Amp | H<sub>2</sub>O | Amp | H<sub>2</sub>O | Amp | H<sub>2</sub>O | Amp |
|  | Incidence | 0/4 (0%) | 4/4 (100%) | 1/4 (0%) | 4/4 (100%) | 4/4 (100%) | 4/4 (100%) | - | - |
| | Burden | <0.1 | $1.52 \times 10^7$ | <0.1 | $1.31 \times 10^7$ | 27 | $1.61 \times 10^7$ | - | - |
|  |  | 1.00E+02 |  | 1.00E+04 |  | 1.00E+06 |  | 1.00E+08 |  |
| CD vs. PD |  | CD | PD | CD | PD | CD | PD | CD | PD |
| Expt A: | Incidence | - | - | - | - | 1/6 (16.7%) | 6/6 (100%) | - | - |
| | Burden | - | - | - | - | <0.1 | $2.58 \times 10^3$ | - | - |
| Expt B: | Incidence | - | - | 2/4 (50%) | 4/4 (100%) | 4/4 (100%) | 4/4 (100%) | 4/4 (100%) | 4/4 (100%) |
| | Burden | - | - | 67 | $2.71 \times 10^4$ | 25 | $2.90 \times 10^4$ | 51 | $3.19 \times 10^4$ |
| Expt C: | Incidence | 0/4 (0%) | 3/4 (75%) | 2/8 (25%) | 5/8 (62.5%) | - | - | - | - |
| | Burden | <0.1 | $3.66 \times 10^2$ | <0.1 | $1.13 \times 10^3$ | - | - | - | - |
| **Total** | **Incidence** | **0/4 (0%)** | **3/4 (75%)** | **4/12 (33.3%)** | **9/12 (75%)** | **5/10 (50%)** | **10/10 (100%)** | **4/4 (100%)** | **4/4 (100%)** |
|  | Chi-squared | p=0.1429 |  | p=0.0995 |  | p=0.0325 |  | P>0.9999 |  |
|  | **Median Peak d7-14** | **<0.1** | **366** | **<0.1** | **1827** | **0.4** | **8912** | **51** | **31923** |

### Persistent CR-Kp colonization is consistently established in a model of pediatric malnutrition

We next investigated whether mice could be colonized without antibiotic-mediated microbiota depletion using an established model of pediatric protein malnutrition (36, 37). Just-weaned juvenile mice (3-4 weeks old) were placed on a complete chow diet (CD, 20% complex protein source) versus an isocaloric purified protein-deficient diet (PD, 2% casein protein) prior to inoculation with Kp175. Typical of this diet, PD-fed mice lost 10-20% of their weight within the first week of diet initiation as compared to CD-fed mice which demonstrated around 50% weight gain (Fig S2A). After 7 days on diet, total bacterial density in fecal samples was similar regardless of diet, and there was no growth of organisms on selective MacConkey with or without meropenem, confirming no Enterobacterales at baseline (Fig S1A,B). Subsequent weight change in mice inoculated with 10^6^-10^8^ CFU Kp175 was similar to mice receiving PBS as a control (Fig S2A) and mice demonstrated normal activity level and stool consistency. PD-fed mice demonstrated a higher burden of fecal shedding at early and later timepoints than CD-fed mice (Fig 2A, Table 1). Intestinal content was cultured from a subset of mice terminated on day 14 demonstrating the greatest burden of colonization in the cecum and colon of PD-fed mice that was >2 logs higher than that of CD-fed mice (Fig 2B). There was no evidence of extra-intestinal spread to the spleen on either diet (Fig 2B).

**Figure 2:**
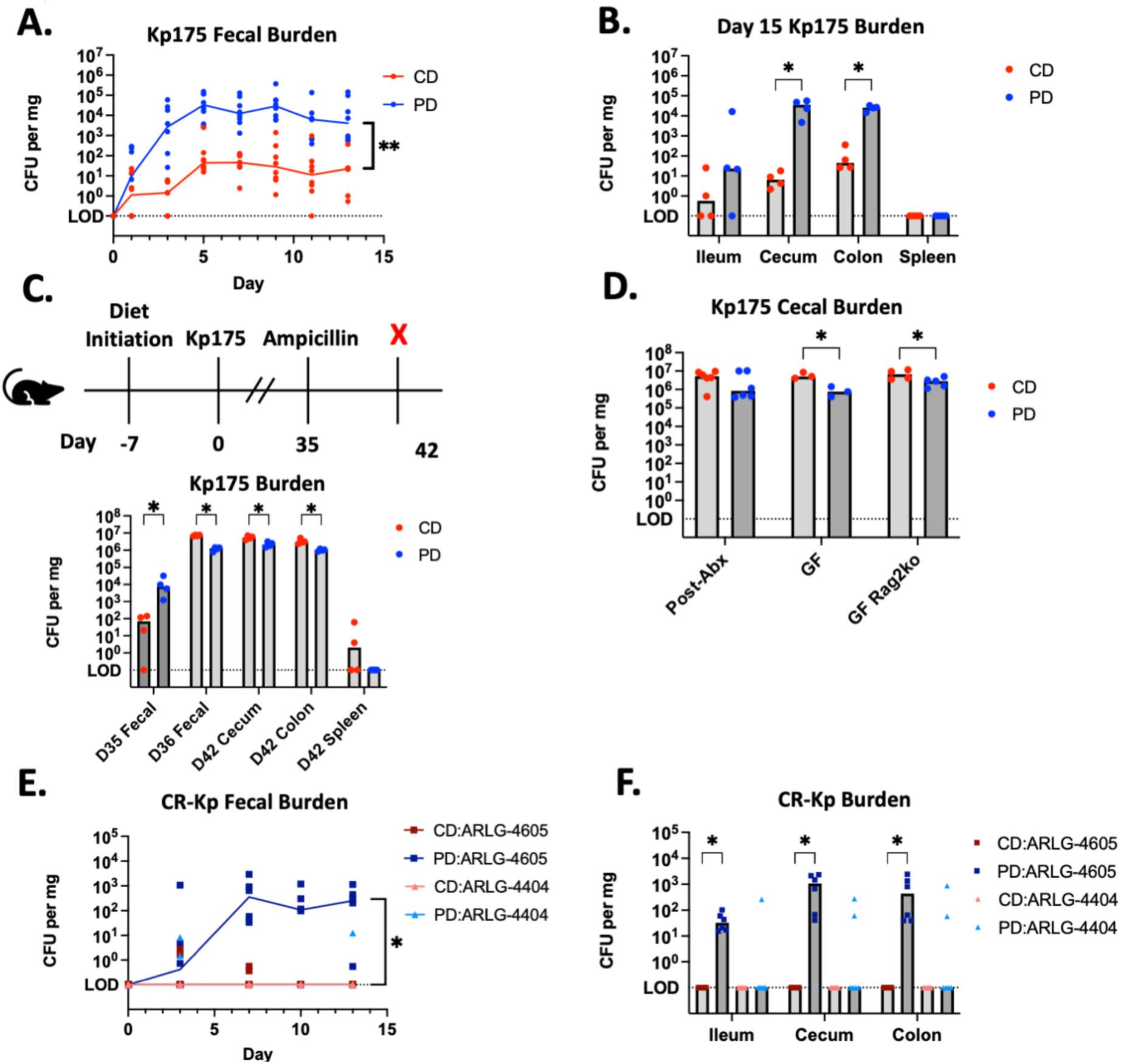
CR-Kp Colonization in Mice on a Protein-deficient Diet. (A.) Mice were initiated on CD or PD 7 days prior to oral gavage with Kp175. Fecal specimens were collected for culture-based quantification at days 0, 1, 3, 5, 7, 9, 11, 13 after Kp175 challenge. Median burden depicted, N=8 per group (4 receiving 10^6^ CFU and 4 receiving 10^8^ CFU on each diet). 2-way ANOVA, **=p<0.01 (B.) Mice originally challenged with 10^8^ CFU terminated on day 15, intestinal contents and spleens harvested for culture-based quantification. Median burden depicted, N=4 per group, Mann-Whitney U-test with 10% FDR, *= p<0.05. (C.) Mice originally challenged with 10^6^ CFU received ampicillin in their drinking water starting on day 35 after challenge with Kp175, terminated on day 42 and intestinal contents and spleens harvested for culture-based quantification. Median burden depicted, N=4 per group, Mann-Whitney U-test with 10% FDR, *= p<0.05. (D.) Mice fed with PD or CD were challenged with 10^6^ CFU Kp175, terminated either 7 days post challenge for Germ-Free wildtype (GF) and GF Rag2^-/-^ mice or 15 days post antibiotic challenge and one day post-ampicillin treatment (Post-Abx) and cecal contents harvested for culture-based quantification. Median burden depicted, N=4 per group, Mann-Whitney U-test with 10% FDR, *= p<0.05. (E.) Mice on CD or PD were challenged with 10^6^ CFU ARLG-4605 or ARLG-4404. Fecal specimens were collected for culture-based quantification at days 0, 3, 7, 10, 13 after challenge. Median burden depicted, N=6 per group. 2-way ANOVA, *=p<0.05 for PD-versus CD-fed mice challenged with ARLG-4605 (F.) CD- or PD-fed mice terminated on day 15, intestinal contents harvested for culture-based quantification. Median burden depicted, N=6 per group, Mann-Whitney U-test with 10% FDR, *= p<0.05.

Remaining mice on both diets demonstrated persistent colonization out to day 35 post-inoculation, with a consistently greater burden in mice fed the PD diet (Fig 2C). In these mice we tested whether disruption of the microbiota would impact colonization dynamics differently depending on diet (Fig 2C). Enteral ampicillin in drinking water on day 35 resulted in an immediate bloom of Kp175 in colonized mice on both diets with a 2-log increase in fecal shedding of Kp175 in PD-fed mice and a 5-log increase in CD-fed mice in feces and intestinal contents (Fig 2C). Overall, there was a slight colonization advantage for Kp175 in feces and intestinal contents of CD-versus PD-fed mice after antibiotic treatment (Figure 2C). These findings suggest a pivotal role for the microbiota, rather than direct effects of the diet or host defense, in determining nutrition-dependent colonization. There was detection of low-level dissemination to the spleen in two CD-fed mice after antibiotic treatment, however there were no signs of systemic illness in these mice (Fig 2C).

A maximal cecal colonization burden of ∼10^7^ CFU/mg in CD-fed mice post-antibiotic treatment was similar to what was observed in the feces of ampicillin pre-treated mice (Fig 1A, Table 1). PD-fed mice had a slightly lower colonization burden of ∼10^6^ CFU/mg (Fig 2E). Germ Free C57BL/6 wildtype and Rag2^-/-^ mice on either diet inoculated with 10^6^-10^8^ CFU Kp175 also achieved roughly equivalent cecal colonization densities as antibiotic-treated mice (Fig 2E). These findings confirm a fixed maximal colonization burden in the absence of microbiota that is not impacted by the adaptive immune system and is slightly reduced by lower dietary protein availability. When the microbiota was left intact, Kp175 colonization of CD-fed mice was inconsistent and dose-dependent with an ID_50_ of 10^6^ CFU and a mean colonization burden of <100 CFU/mg in the feces regardless of dose (Table 1). In contrast, Kp175 consistently colonized PD-fed mice with an ID_50_ of ≤10^2^ CFU and at a fecal burden 3-4 logs higher than observed in CD-fed mice (Table 1). Thus, dietary protein-deficiency is sufficient to impart a colonization burden that is intermediate to colonization achieved with depletion or removal of the microbiota.

We additionally investigated whether protein-deficiency supported colonization of other clinical isolates of CR-Kp (Table S1). After inoculation with 10^6^ CFU, ARLG-4605 achieved colonization in feces and intestinal contents of PD-fed mice similar to Kp175, however it did not colonize CD-fed mice (Fig 2E,F). In contrast, ARLG-4404 was incapable of colonizing mice regardless of diet (Fig 2E,F). These findings demonstrate variable nutrition-dependent colonization across select CR-Kp strains.

### Persistent colonization in PD-fed mice is not reversed with dietary rehabilitation

Based on the observation that diet determines CR-Kp colonization, we hypothesized that nutritional repletion would reverse colonization in PD-fed mice. Kp175 colonization was established in a cohort of PD-fed mice and then half of these mice were transitioned to CD (Fig 3A). Although the mice switched to CD demonstrated catch-up growth (Fig S3B), the diet change failed to significantly reduce Kp175 colonization burden with persistent shedding seen beyond 40 days at burdens similar to mice that remained on PD (Figure 3A). These results demonstrate that once established, nutrition-dependent colonization with CR-Kp is long lasting both in mice that remain malnourished and in those that have returned to a healthy nutritional status.

**Figure 3:**
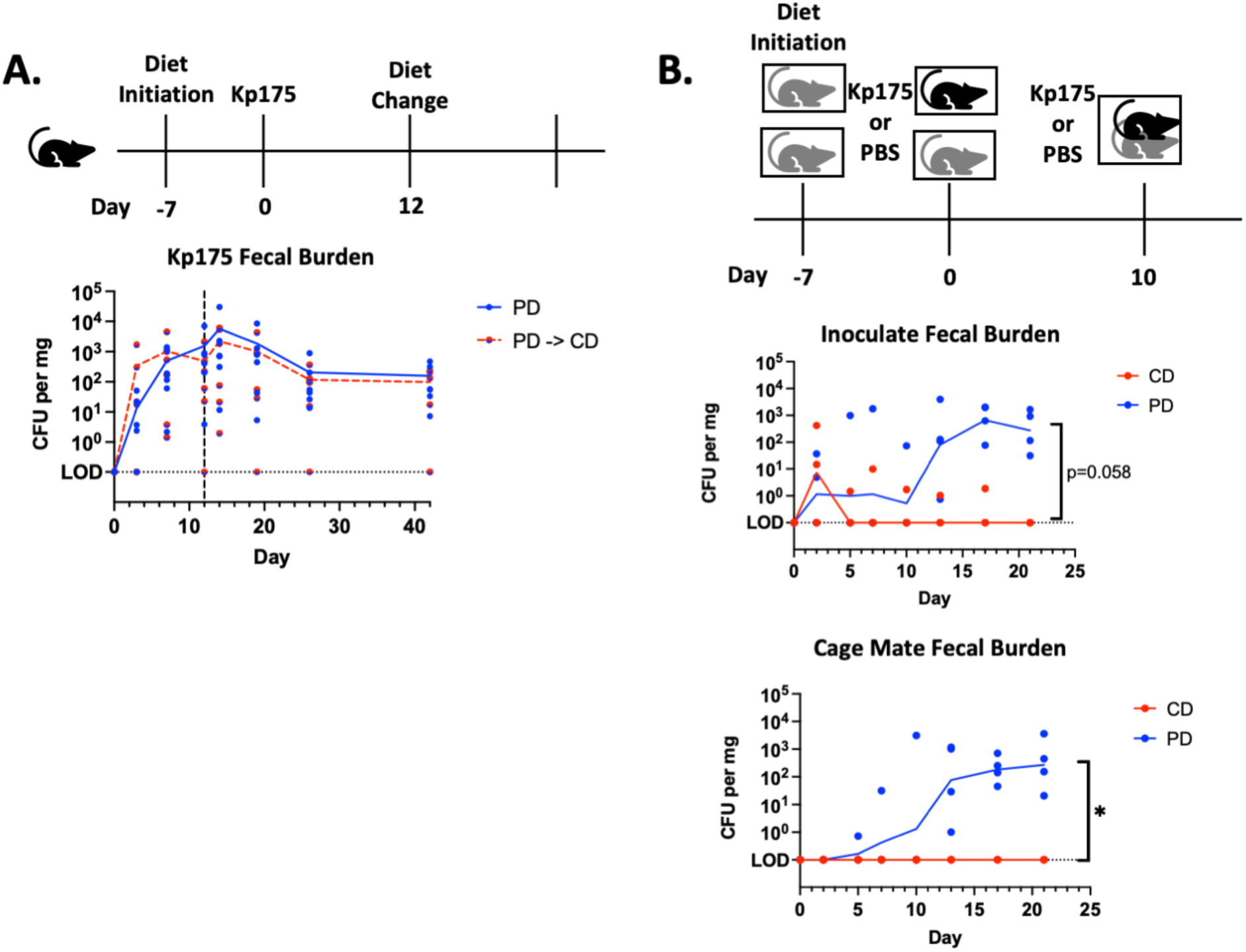
Kp175 Colonization After Refeeding and Transmission after Co-housing. (A.) Mice initiated on PD 7 days prior to oral gavage with 10^6^ CFU Kp175. On day 12, half of mice were switched from PD to CD and the other half were continued on PD. Fecal specimens were collected for culture-based quantification on days 0, 7, 10, 12, 14, 19, 28, and 42 after challenge. Median burden depicted, N=8 per group. (B.) Mice initiated on respective diets 7 days prior to oral gavage with 10^6^ CFU Kp175 or PBS, with one Kp175 inoculated mouse, and one PBS control cage-mate placed together in a single cage on day 2. A subsequent challenge with 10^6^ CFU Kp175 was given to the originally inoculated mouse on day 10 if colonization had not been established (N=3 CD fed mice, and N=0 PD fed mice). Fecal specimens were collected for culture-based quantification on days 0, 2, 5, 7, 10, 12, 17, 21 after challenge. Median burden depicted, N=4 per group. 2-way ANOVA, *=p<0.05.

### Dietary protein deficiency permits host-to-host transmission of Kp175

To test the impact of persistent colonization in PD-fed mice, transmission experiments were performed in which mice were initially singly housed on their respective diets. After inoculation with Kp175, mice were co-housed with a PBS-inoculated cage-mate on the same diet (Figure 3B). Transmission was observed between all Kp175 inoculated PD-fed mice and their unexposed cage-mates, but not between Kp175 inoculated CD-fed mice and their respective cage-mates, even after a second inoculating dose was given to the colonized cage mate (Figure 3B). These findings demonstrate that PD supports elevated colonization burden and transmission of CR-Kp from a colonized host to its cage-mate. In contrast, transmission does not occur in CD-fed mice.

Since the custom PD diet is also processed with different components than the CD diet, we endeavored to determine if differences in colonization and transmission were due to reduced protein content of PD in comparison to a control purified protein diet. Mice fed an isocaloric custom complete-protein diet (CP, 20% casein protein) also demonstrated lower colonization and transmission than the PD-fed mice (Figure S3). This confirmed that inadequate protein had a specific effect on susceptibility to colonization and transmission beyond other component differences between the PD and CD diets.

### Clinical CR-Kp isolates have variable in vitro growth in protein-deficient conditions

To directly determine how nutritional components might variably support CR-Kp growth, different isolates were grown under various conditions *in vitro*. All tested strains had robust growth in nutritionally rich Brain Heart Infusion media and no growth in M9 minimal media lacking a nutrient source (Fig 4A). To further characterize nutrients that determine *in vitro* growth, CR-Kp isolates were grown in M9 supplemented with glucose as a carbohydrate energy source and/or casein peptone as a protein energy source (Fig 4A). Minimal growth was observed for all isolates with casein peptone alone, and variable growth was observed with glucose alone, with growth of Kp175 > ARLG-4605 and no growth of ARLG-4404 (Figure 4A). It was only with supplementation with both glucose and casein peptone that all 3 isolates demonstrated robust growth (Fig 4A). These findings demonstrate isolate-dependent nutrient requirements for growth, with Kp175 growing better than other strains in the absence of protein and ARLG-4404 requiring both a carbohydrate and protein/amino acid source to grow.

**Figure 4:**
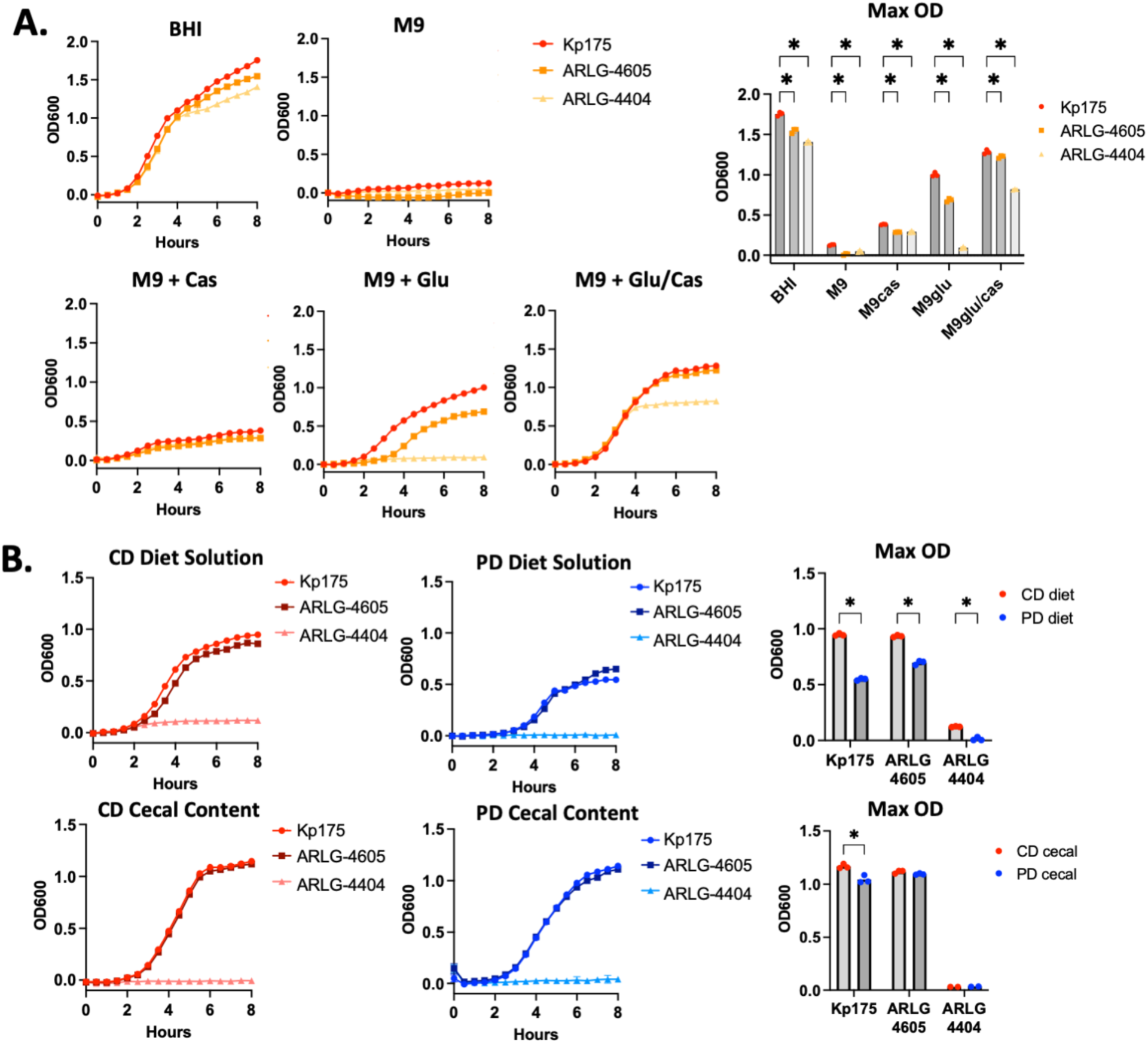
CR-Kp Growth in Nutrient-Limited Conditions in vitro. (A.) Density as determined by OD600 of CR-Kp strains in BHI, plain M9 or M9 supplemented with glucose (4000 ug/mL) and/ or casein peptone (1600 ug/mL) as indicated. Mean Density depicted, N=3 for each condition plotted. Mann-Whitney U test with 10% FDR comparing peak OD600, *= p<0.05. (B.) M9 + glucose (4000 ug/mL) supplemented with 1% by weight CD or PD food or filtered CD or PD cecal content normalized to 160 ug/mL protein content. Mean Density depicted, N=3 for each condition plotted. Mann-Whitney U test with 10% FDR comparing peak OD600, *= p<0.05.

We further investigated how the nutrients available from the diet itself might impact the growth of CR-Kp strains differently *in vitro.* Kp175 and ARLG-4605 demonstrated robust growth in media-derived from CD (1% diet in M9/glucose) with reduced growth in media derived from PD (Fig 4B). No growth was seen in either dietary solution for ARLG-4404 (Fig 4B). In contrast, robust growth of Kp175 and ARLG-4605 was observed in media supplemented with cecal content supernatants from both CD-fed and PD-fed mice (160 ug/mL of cecal content protein in M9/glucose), with only a slight growth advantage for Kp175 in CD versus PD cecal content supplemented media (Fig 4B). No growth was seen in either cecal content supplemented media solution for ARLG-4404 (Fig 4B). These findings demonstrate nutritional components derived from both diets (before and after digestion) adequately support growth of certain CR-Kp isolates, and that in the absence of competing microbes,CD diet components promote growth more effectively than PD diet components supporting observations in ampicillin-treated mice (Fig 2C). Furthermore, the inability of ARLG-4404 to grow in either CD- and PD-supplemented conditions or in the absence of a protein source suggests that strain variability in nutrient utilization may determine nutrition-dependent colonization variability observed *in vivo*.

### Free D- and L-amino acid content in the cecum is not impacted by dietary protein deficiency and does not correspond to colonization susceptibility

LC-MS of the diets confirmed reduced free L-amino acid (microbiota, diet, and/or host-derived) and D-amino acid (diet and/or microbiota-derived) content in PD food relative to CD food (Fig S4A). This could explain the reduced *in vitro* growth observed in PD versus CD-supplemented media (Fig 4B). In contrast, cecal contents of mice fed CD versus PD showed very similar free amino acid availability (Fig S4B,C). This could explain relatively similar *in vitro* growth of Kp175 and ARLG-4605 in media supplemented with CD and PD cecal content supernatants (Fig 4B). In comparison to naïve mice, Kp175 colonization on either diet did not alter free cecal amino acid content. In contrast, there was a significant reduction in nearly all D- and L-amino acids in CR-Kp colonized mice on either diet after ampicillin treatment (Fig S4B,D). Collectively these findings suggest that protein utilization requirements vary across different CR-Kp strains, with certain strains such as Kp175 able to thrive *in vitro* and *in vivo* with minimal protein/amino acid availability. Furthermore, cecal amino acid availability does not appear to limit colonization.

### Bile acid profiles correlate with CR-Kp colonization burden

Since amino acid availability was not the factor determining nutrition-dependent colonization in CD-versus PD-fed mice, we next investigated whether PD led to loss of microbiota-derived molecules that may inhibit Kp colonization. Secondary bile acids are prototypic metabolites that are regulated by the microbiota with variable composition dependent on dietary protein availability (38–40). LC-MS revealed that ampicillin, regardless of diet, led to expected reduction in many unconjugated primary and secondary bile acids, consistent with the known role of the microbiota in bile acid deconjugation and secondary bile acid generation (Fig 5A,B) (41).

**Figure 5:**
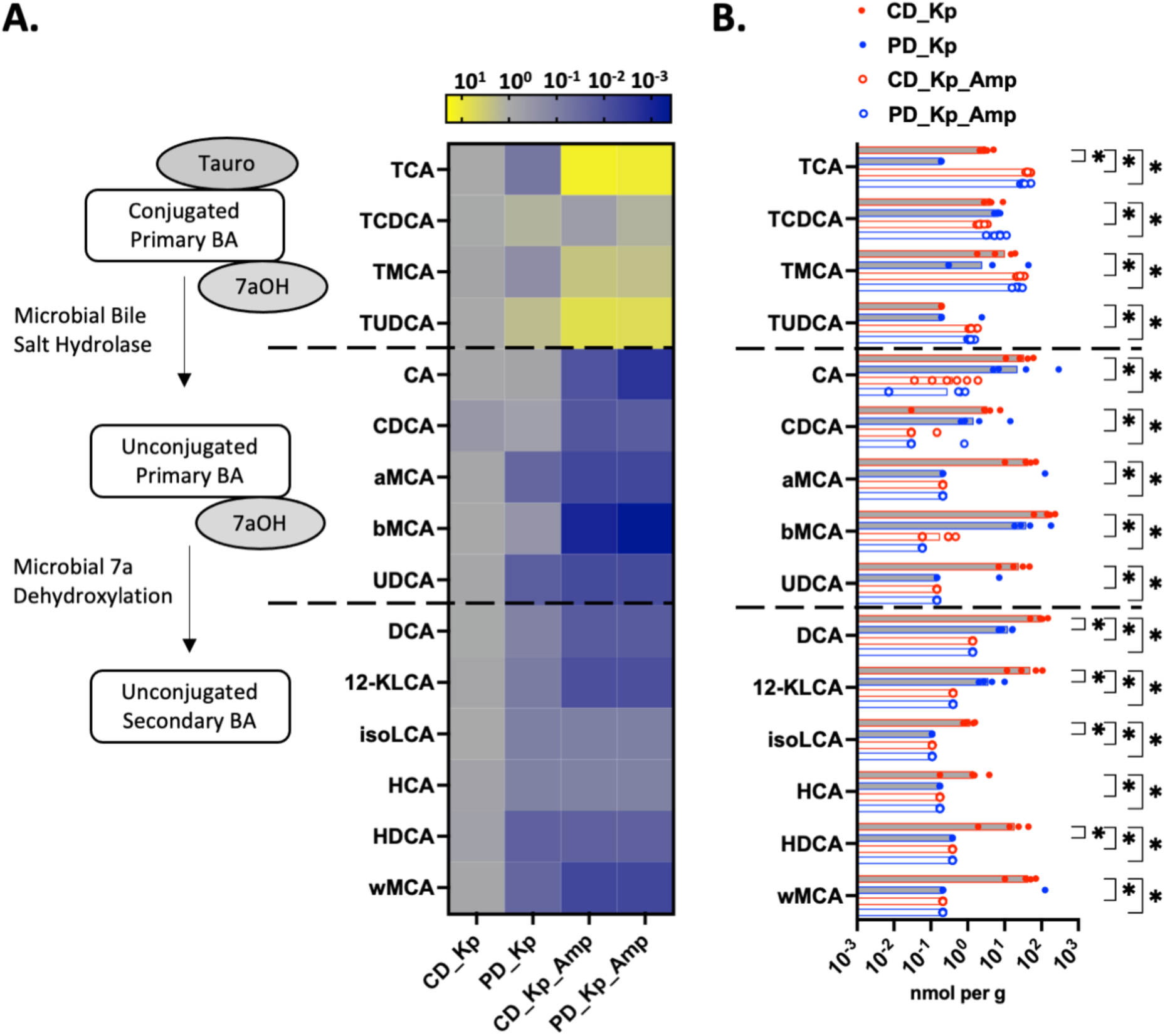
Bile Acid Content in Ceca of Colonized Mice With or Without Ampicillin Treatment. (A.) Heat map of individual bile acids in cecal content of mice placed on respective diets 7 days prior to inoculation with 10^6^ CFU and terminated on day 14 post-inoculation or treated with Ampicillin in the drinking water (1 mg/ml) at day 35 post-inoculation and terminated at day 42. Expressed fold change relative to average concentration in cecal contents of colonized CD-fed mice without antibiotic treatment. Median concentration depicted, N=4-6 per group. (B.) Concentration of individual bile acids in cecal content of mice placed on respective diets 7 days prior to inoculation with 10^6^ CFU and terminated on day 14 post-inoculation or treated with Ampicillin in the drinking water (1 mg/ml) at day 35 post-inoculation and terminated at day 42. Median concentration depicted, N=4-6 per group. Mann-Whitney U with 10% FDR test for each bile acid, *= p<0.05 for comparison of CD-fed mice without antibiotics to other groups. *TCA= Taurocholic Acid, TCDCA= Taurochenodeoxycholic Acid, TMCA= Tauromuricholic Acid, TUDCA= Tauroursodeoxycholic Acid, CA= Cholic Acid, CDCA= Chenodeoxycholic Acid, aMCA= Alpha Muricholic Acid, bMCA= Beta Muricholic Acid, UDCA= Ursodeoxycholic Acid, DCA= Deoxycholic Acid, 12-KLCA= 12-Ketolithocholic Acid, isoLCA= Isolithocholic Acid, HCA= Hyocholic Acid, HDCA= Hyodeoxycholic Acid, wMCA= Omega Muricholic Acid*.

Consistent with our prior findings, there was a statistically significant reduction in unconjugated secondary but not primary bile acids in PD-vs CD-fed mice in the absence of antibiotic treatment (Fig 5B) (38). Colonization with Kp175 alone did not significantly alter bile acid profiles on either diet (Fig S5). When comparing bile acid content and Kp175 burden in cecal contents of colonized mice, there was a statistically significant inverse correlation between colonization burden and quantity of individual unconjugated bile acids and total primary and secondary unconjugated bile acids (Fig 6A, Table S2). In comparison, there was a positive correlation between colonization burden and quantity of certain individual conjugated primary bile acids and total conjugated primary bile acids (Fig 6A, Table S2). These findings demonstrate that lower concentrations of unconjugated bile acids, particularly secondary bile acids, in mice fed PD and/or treated with antibiotics correspond with higher burdens of colonizing Kp175.

**Figure 6:**
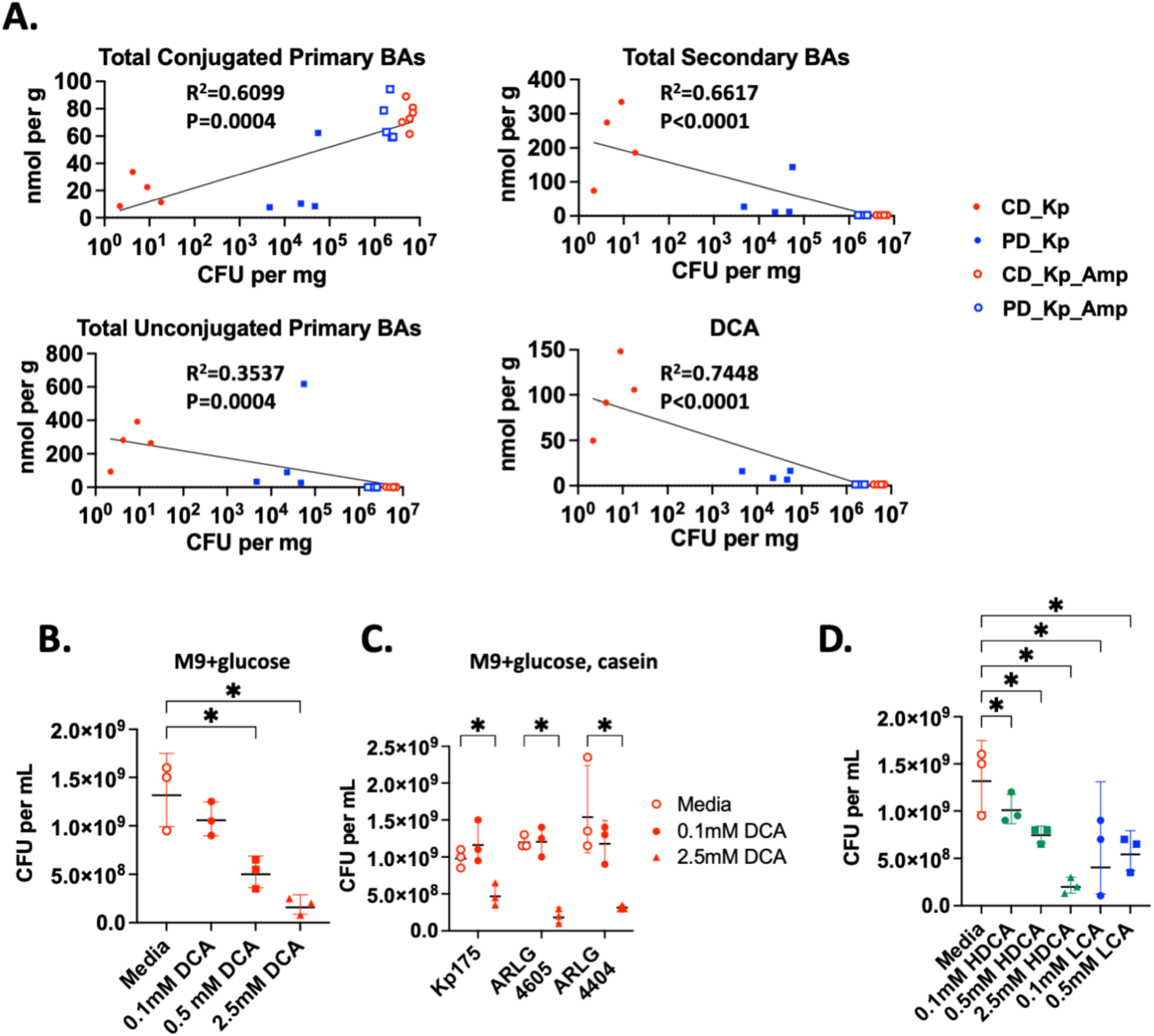
CR-Kp Growth in Media Supplemented with Secondary Bile Acids. (A.) Concentration of total conjugated primary bile acids, total unconjugated primary bile acids total secondary bile acids, and deoxycholic acid (DCA) plotted against Kp175 burden in ceca of mice. Line of best fit and R^2^ calculated with least squares regression with X as logarithmic variable and Y as linear variable and p calculated using Spearman correlation method. (B.) Quantity of Kp175 recovered on BHI agar after 5×10^6^ CFU per mg seeded in M9 + glucose (4000 ug/mL) 2.5% DMSO and Deoxycholic Acid (DCA) supplemented at concentrations indicated and grown for 24 hr. Mean concentration depicted, N=3 per condition. Mann-Whitney U with 10% FDR test for media vs. other conditions. (C.) Quantity of CR-Kp recovered on BHI agar (mean +/-SEM) after 5×10^6^ CFU per mg seeded in M9 + glucose (4000 ug/mL) and casein peptone (1600 ug/mL) with 2.5% DMSO and DCA supplemented at concentrations indicated. Mean concentration depicted, N=3 per condition. Mann-Whitney U with 10% FDR test for media vs. other conditions. (D.) Quantity of Kp175 recovered on BHI agar (mean +/-SEM) after 5×10^6^ CFU per mg seeded in M9 + glucose (4000 ug/mL) and casein peptone (1600 ug/mL) with 2.5% DMSO and Hyodeoxycholic Acid (HDCA) or Lithocholic Acid (LCA) supplemented at concentrations indicated. Mean concentration depicted, N=3 per condition. Mann-Whitney U with 10% FDR test for media vs. other conditions.

### Secondary bile acids inhibit growth of CR-Kp in vitro

Among individual bile acids, the strongest negative correlation with Kp175 burden was observed with deoxycholic acid (DCA), one of the most plentiful secondary bile acids in our mice and in humans (Fig 6A, Table S2) (39). We therefore hypothesized that DCA might directly inhibit CR-Kp growth. We observed that DCA caused dose-dependent inhibition of Kp175 that was present in media supplemented with protein and glucose and most pronounced in media with only glucose (Fig 6B,C). Dose dependent inhibition by DCA was additionally demonstrated for ARLG-4404 and ARLG-4605 in protein-replete conditions (Figure 6C). Other secondary bile acids including hyodeoxycholic acid (HDCA) and lithocholic acid (LCA) also demonstrated dose-dependent inhibition of Kp175 growth *in vitro* (Figure 6D). Collectively these findings confirm that secondary bile acids inhibit growth of a range of CR-Kp isolates under various conditions and may be implicated in nutrition-dependent colonization.

## Discussion

Using a mouse model of pediatric protein malnutrition and clinical isolates of CR-Kp we have demonstrated nutrition-dependent intestinal colonization resistance against HRE. Our results show that juvenile mice fed a protein-deficient diet are susceptible to establishment of intestinal colonization with specific strains of CR-Kp at higher burdens than experienced in mice fed a complete nourished diet. This nutrition-dependent colonization burden is sufficient to allow for transmission between hosts and can be established with an inoculating dose as low as 100 CFU, well below the estimated quantity of Enterobacterales ingested by children in certain community environments in low- and middle-income countries (42). We further observed that HRE colonization established in malnourished hosts cannot be reversed by dietary rehabilitation. These findings have important implications for the establishment and persistence of HRE in susceptible global populations. Protein malnutrition is most commonly encountered in children and elderly adults residing in low- and middle-income countries in Sub-Saharan Africa and South Asia, regions of the world that also experience some of the highest rates of infection and mortality from HRE including CR-Kp (1, 2, 12, 13, 43, 44). Malnourished individuals across the globe have increased exposure to covariate risks for HRE colonization and infection including healthcare contact, antibiotic use, and co-morbidity burden (45, 46). This study establishes a direct causal link between inadequate protein intake itself and HRE colonization.

Several publications have investigated how dietary alteration can impact intestinal colonization by Enterobacterales in animal models. A high fiber diet was shown to reverse *K. pneumoniae* colonization *in vivo* by promoting a diverse microbiota capable of restricting simple carbohydrate availability (47). In contrast, a low fiber diet promoted pathogenesis of the model colitogenic pathogen *Citrobacter rodentium* through microbiota-mediated mucosal digestion for carbohydrate acquisition (48). Common to these studies is the importance of nutrient availability in determining Enterobacterales colonization. Multiple recent studies have reinforced that antibiotic-mediated depletion of the microbiota promotes Enterobacterales colonization by eliminating competition for specific nutrients, particularly certain simple carbohydrates and amino acids (22, 49). We similarly hypothesized that alteration in nutrient availability on a protein-deficient diet might impact diet-dependent colonization by CR-Kp.

Indeed, we observed variable growth of CR-Kp isolates *in vitro* in protein-limited conditions, with our poorest colonizing isolate, ARLG-4404, unable to grow without a protein/amino acid source. In contrast, Kp175 and ARLG-4605 demonstrated robust growth in the absence of protein *in vitro*. Thus, certain CR-Kp strains may thrive in protein-poor conditions *in vivo* that might hinder other intestinal commensals competing for nutrient energy sources. However, in our model we found that free amino acid composition in the ceca of colonized mice was largely unaffected by dietary protein input. In contrast, amino acids were reduced in mice experiencing high colonization burdens post-antibiotic treatment on either diet. These findings are consistent with microbiota-mediated engineering of the intestinal microenvironment and indicate that amino acid availability is unlikely to be the factor determining HRE colonization in our model.

Previous work utilizing this pediatric protein malnutrition model demonstrated that the presence of an intact microbiota was necessary for mediating nutrition-dependent colonization burden and disease pathogenesis during experimental infection with diarrheagenic protozoans and Enterobacterales (29, 32, 35–37). Indeed, substantial changes in the 16S composition of the microbiome have been demonstrated on a protein-deficient diet with corresponding changes in certain key metabolites (29, 33). In particular, we recently demonstrated that there is alteration in the bile acid composition in mice on a protein-deficient diet with an increased ratio of primary to secondary and conjugated to unconjugated bile acids (38). In the current study we again found a reduction in secondary bile acids during protein deprivation that correlates significantly with increased colonization with CR-Kp. Another recent study showed reductions in secondary bile acid content in mice rendered susceptible to intestinal colonization with *K. pneumoniae* through alcohol-induced dysbiosis (50). In this study, investigators observed that the secondary bile acid LCA restricts *K. pneumoniae* growth and binding to epithelial cell lines *in vitro*. We similarly found that multiple secondary bile acids restricted CR-Kp growth *in vitro*, suggesting a possible direct impact of secondary bile acids in restricting intestinal colonization *in vivo*. There is limited data regarding secondary bile acids in the lower intestinal tract of malnourished versus healthy children (51). However, several studies have shown increased conjugated primary bile acids in the serum of acutely and chronically malnourished children, similar to what we observed in cecal contents of PD-fed mice (51, 52). Collectively, these studies suggest that regulation of bile acids during pediatric malnutrition demands further characterization, particularly in the large intestine where these metabolites may impact colonizing pathogens including *Klebsiella* and other Enterobacterales.

There is substantial clinical literature to support the importance of dietary nutrition on infection with a variety of intestinal-dwelling bacteria (53). Malnutrition is associated with increased carriage and infection with diarrheagenic Enterobacterales in children in Sub-Saharan Africa and South Asia (54). Pediatric malnutrition is also associated with onset of bacteremia from “opportunistic” intestinal colonizing Enterobacterales including *Klebsiella, Salmonella* and non-toxigenic *E. coli* (15, 55). Few studies to date have specifically investigated HRE colonization and infection in malnourished individuals, however extremely high rates of colonization were found in children admitted with severe acute malnutrition in a hospital in Niger, and malnutrition was identified as an independent risk factor for HRE bloodstream infections in children in Senegal (13, 14, 17). In a cohort in Malawi, we recently found 80% HRE colonization in children upon hospital admission for acute malnutrition (56). Our results here support the hypothesis that malnourished hosts have an independent biological predisposition to intestinal colonization with Enterobacterales including HRE. Also notable is our finding that only some isolates are capable of colonizing protein-deficient hosts, suggesting that certain strains may pose a greater risk for spread through malnourished populations. These findings are of particular importance to the global proliferation and spread of HRE. International guidelines currently recommend that children diagnosed with severe acute malnutrition receive empiric treatment with a broad beta-lactam antibiotic such as amoxicillin (19). Increased likelihood of colonization with HRE organisms resistant to such empiric therapy must be considered to avoid promoting resistant infection and transmission in malnourished children undergoing guideline-directed treatment.

While our model of protein malnutrition recapitulates key characteristics of human pediatric stunting from conventional intestinal pathogens, there is a vast spectrum of human malnutrition across global space and the lifespan that may function in different ways to alter HRE colonization risk. Further work using dietary formulations that more closely model human dietary intake and/or address different macro- and micronutrient deficiencies will be necessary to fully elucidate the complicated relationship between dietary nutrition and HRE colonization. Additional work in these models should investigate how individual members of the microbiota might play a role in nutrition-dependent colonization resistance, and how bile acids and other metabolites might serve as putative HRE inhibitors. In light of the heterogeneity seen among our selection of only three different CR-Kp isolates, future studies should also utilize a variety of strains and species of Enterobacterales resistant or susceptible to carbapenems and/or 3^rd^ generation cephalosporins.

Much remains to be investigated and discovered related to nutrition-dependent colonization resistance. However our current results are novel and important in several regards. Here we demonstrate a direct causal link between dietary malnutrition and colonization with opportunistic antimicrobial resistant pathogens. These findings also reveal the importance of the microbiota in mediating protein malnutrition-dependent colonization. In particular they suggest a role for microbiota-derived secondary bile acids which have remained largely unexplored in their relationship to HRE growth and persistence. Finally we describe here a novel animal model system that permits colonization with HRE through dietary alterations alone without antibiotic-mediated microbiota depletion that is required in most intestinal colonization models (57). We expect this model will serve as the basis of future work investigating how nuanced manipulation of the microbiota without antibiotic treatment impacts HRE colonization. The work described here and the studies to follow will help to build a detailed mechanistic understanding of how diet regulates HRE colonization, and how this might be intervened on through nutritional and probiotic approaches to address the devastating global impact of these pathogens.

## Methods

### Sex as a Biological Variable

Prior investigations in our lab have found no impact of sex in mouse models of diet-dependent colonization and infection with enteric pathogens, and have generated metabolomic and microbiome data in male mice on a protein-deficient diet (29, 35, 38). Thus, in this study we utilized male mice to minimize variability in our phenotype while using the least number of animals necessary according to the protocol approved by our Institutional Animal Care and Use Committee (IACUC).

### Bacterial Isolates and Growth

*Klebsiella pneumoniae* isolates used in this study were originally obtained from human specimens as reported in Table 1. Bacteria from frozen 20% glycerol stocks were streaked onto Brain Heart Infusion Agar (BHI, BD) and incubated overnight at 37°C, then individual colonies were picked with a sterile loop and inoculated into BHI broth and incubated with constant agitation at 37°C to log phase, 1.5-2 hr. Bacteria were centrifuged 10 min, 3000xg, 4°C and pellets washed with sterile Dulbecco’s Phosphate-Buffered Saline (DPBS, Gibco). Density was estimated with a Spectrophotometer (Bio-Rad Smart-Spec Plus) and bacteria were resuspended to desired concentration in sterile DPBS. Final concentration was determined by performing serial dilution in sterile DPBS to 10^-6^ and spreading 10 uL per quadrant in duplicate onto BHI agar plates. After incubation at 37°C overnight colonies were counted and colony forming units (CFUs) per mL were calculated. Criteria for strain selection included overlapping growth curves in BHI during continuous monitoring and similar recovery from MacConkey agar containing selective antibiotics.

### Animal Models and Handling

C57BL/6 male mice were acquired from Jackson Laboratories. For experiments with mice receiving regular drinking water or ampicillin in their water, adult mice 6-12 weeks for used. For experiments comparing animals on different diets, mice were obtained immediately after weaning at age 3-4 weeks and weighing 10-15 grams each. Mice were housed in the UNC National Gnotobiotic Rodent Resource Center (NGRRC)under specific pathogen free conditions. Alternatively, where indicated, Germ Free C57BL6 wildtype and Rag2^-/-^ mice of either sex bred in the UNC NGRRC were housed under Germ Free conditions and used at any available age (12-28 weeks), with matching between groups for age, weight, and sex prior to initiation of experiment. According to the approved IACUC protocol, animals were co-housed at 2 per cage to minimize stress on animals on nutrient-deficient diets. Exceptions were made for mice housed in Germ Free conditions, and in transmission experiments in which mice were temporarily placed into individual cages on the day of inoculation, then selectively co-housed at 2 per cage on day 2 after inoculation. Cages, water, and food were autoclaved, and sterile dry-rite bedding was used to minimize microbial contamination. Mice were fed a fully-nourished, complex chow-based diet (CD) with 20% protein by weight (Teklad Global Soy Protein-Free Extruded Rodent Diet 2020SX, Inotiv), a custom isocaloric purified casein protein-deficient diet (PD) with 2% protein by weight (TD.110200, Inotiv), or a custom isocaloric protein-adequate purified casein protein diet (CP) with 20% protein by weight (TD.08678, Inotiv). Mice were handled using gnotobiotic techniques in all experiments to eliminate microbial contamination and between-cage transmission, with additional sterilization with bleach and peroxide solution and fresh sterile gloves used between each group of animals in each experiment. Mice were weighed every other day with a digital battery-operated scale (Ohaus) with each mouse placed in a separate autoclaved plastic box prior to weighing.

### Animal Model Colonization

Prior to inoculation with bacteria, mice were allowed to equilibrate to the facility on their respective diet for at least 1 week. In select experiments, normal facility drinking water was replaced with facility water with 1 mg/mL of ampicillin (GoldBio) either before or after inoculation and mice were allowed to drink *ad libitum*. Bacterial isolates were grown as described above and resuspended in sterile DPBS at 10^3^-10^9^ CFU per mL and kept on ice until inoculation. Orogastric gavage was performed with 100 µL of (10^2^-10^8^ CFU) per mouse. Fresh fecal pellets were collected into sterile pre-weighed Eppendorf tubes at set time points. Tubes were weighed and fecal pellets were subsequently resuspended in 250 µL sterile DPBS and homogenized with a Motorized Tissue Grinder (Fisher) prior to performing serial dilution to 10^-6^ in sterile DPBS. 10 µL of dilution were spread per quadrant in duplicate onto MacConkey agar (BD) plates containing 125 µg per mL Meropenem (GoldBio). After incubation at 37°C overnight colonies were counted and CFUs per mL were calculated. Limit of Detection (LOD) was set at 0.1 CFU/mg (corresponding to growth of just 1 colony from a 25 mg fecal pellet, the smallest detectable with our culture method). At the end of each experiment mice were euthanized via CO_2_ inhalation and intestinal tissue along with spleen were collected. Intestinal contents were pushed out and specimens were weighed, homogenized, and cultured for bacterial quantification as described above. Routine culture of fecal pellets on MacConkey with Meropenem was also performed prior to bacterial gavage in all experiments to confirm no prior colonization with carbapenem-resistant Enterobacterales.

### Quantitative PCR (qPCR)-based quantification of total bacteria

DNA was purified from log-phase Kp175 grown in culture in BHI using a Qiagen Dneasy Blood and Tissue Kit and from mouse fecal pellets using a Qiagen QIAamp Fast DNA Stool Kit per manufacturer’s instructions. Quantification of total 23S rRNA was performed as previously described (58). PCR reactions were prepared with 25 uL per well in a 96 well qPCR plate consisting of: 12.5 uL 2x Bio-Rad iQ Supermix, 1 uL each of forward and reverse primer, 0.5 uL probe, 5 uL Microbial DNA-free water and 5 uL template DNA. DNA from Kp175 grown in culture was used as a standard. Oligonucleotide sequences for primers were ATTACGCCATTCGTGCAGGTCGGA for 23S forward, TAAACGGCGGCCGTAACTATAACGGT for 23S reverse, and TAMRA-CCTAAGGTAGCGAAATTCCTTGT-MGB for 23S probe. qPCR was run on a QuantStudio 3 Real Time PCR System (Thermo Fisher) with the following protocol: 50°C x 2min, 95°C x 15min, [94°C x 1min, 60°C x 1min] x 40 cycles.

### In Vitro Growth Assays

Bacteria were grown to log phase as described above and pelleted 10 min, 3000xg, 4°C and washed with 10 mL M9 base (Gibco M9 salt solution, 2 mM MgSO_4_, 0.1 mM CaCl_2_) prior to resuspending in M9 base at OD600 of 0.1 as determined by spectrometer. Bacteria were then diluted 1:10 in their respective growth media for growth assays. Media used in these experiments included BHI and M9 base supplemented with L-glucose (Fisher), casein peptone (Thermo-Fisher) or diet solution or cecal solutions. Diet solutions were prepared by grinding CD and PD pellets with a mortar and pestle then adding to M9 base at 10% weight-by-volume and vortexing prior to passing through a 40 µM cell strainer followed by a 0.2 µM syringe filter. Cecal solutions were prepared by pelleting cecal contents of CD- or PD-fed mice (see above) 10 min, 3000xg, 4°C and passing supernatant through a 0.2 µM syringe filter. Protein content of diet and cecal solutions was determined using BCA assay (Thermo-Fisher), and solutions were further diluted as indicated in M9 base. Growth assays were performed in 96 well plates with 200 µL of bacteria dilutions per well in triplicate with serial OD600 measurements read on a Clariostar Plus plate reader (BMG Labtech) over 8 hours of incubation at 37°C. Maximum OD values were determined for each strain grown under the same conditions Alternatively, secondary bile acids including deoxycholic acid, hyodeoxycholic acid and lithocholic acid (Sigma) were dissolved at 100 mM in DMSO prior to addition at desired concentrations to bacterial suspensions in M9 with indicated nutrients and total DMSO of 2.5% in all conditions. Bacteria treated with bile acids and/or DMSO were grown for 24 hours at 37°C. in 96 well plates with 200 µL of bacteria dilutions per well in triplicate and CFU per mL were determined by culturing on BHI agar as described above.

### Quantification of Cecal Amino Acid and Bile Acid Content

Murine cecal content was homogenized with 1 mL of LC-MS grade water using a FastPrep-24™ 24onization24 (2 cycles, 20 seconds). Homogenized samples were centrifuged for 15 minutes at 13,000 x *g* at 4°C, and the supernatant was transferred to a new microtube. A biological quality control (BQC) was prepared by pooling 50 μl aliquots of each study sample.

To prepare the calibration curves of the bile acids, individual stock solutions of 57 target bile acids (final concentration of 10 μM) and 16 internal standards (50 μM) were prepared in H2O:I:IPA (2:1:1, v/v). Mixed calibration standard solutions containing all 57 bile acids were prepared with H2IACN: IPA (2:1:1, v/v) as the solvent, including the following concentrations 5 nM, 10 nM, 25 nM, 50 nM, 100 nM, 250 nM, 500 nM, 1000 nM, 2000 nM, 4000 nM, 6000 nM, 8000 nM, and 10000 nM. Quality control (QCs) samples were prepared for the lowest concentration (QC1, 25 nM), intermediary (QC2, 1000 nM), and highest concentration (QC3, 10000 nM). The internal standard stock solution was used to prepare the internal standard working solution (2.1 μM final concentration).

A 20x dilution was performed for all the samples and BQC for bile acid extraction. Then, 50 µl of diluted sample was transferred to a 2 ml microtube, followed by 10 µl internal standard working solution, and 150 μl of cold methanol. A double blank (diluent only, without the internal standard working solution) and single blank (diluent with 10 µL of the internal standard working solution) was prepared to measure background noise. Samples were vortexed, centrifuged for 15 minutes at 4°C and 13,000 x *g* and the supernatant (100 µl) was transferred to a 96-well plate.

Chromatographic separation was performed on a Waters Acquity UPLC XEVO TQ-XS mass spectrometer equipped with an electro-ionization (ESI) source. The optimal separation was achieved on column ACQUITY BEH C8 column (1.7 μm, 100 mm × 2.1 mm), mobile phase A (2L of UHPLC-UV grade water, 200 ml of acetronitrile, 0.17g ammonium acetate, and 700 µl acetic–acid - final pH 4.16), and mobile phase B (acetonitrile and 2-propanol, 1:1). The gradient separation was completed in 18 minutes with the initial conditions of 90 % of A and flow rate of 0.6 ml/min for 0.1 min. Then, solvent B was increased to 35 %, 65 %, and 100 % over 13.1 minutes at a flow rate of 0.6-1 ml/min. Mobile phase B decreased to 60 % and 40% for 0.2 min until return to the initial condition (90 % A at 0.6 ml/min) for 5 minutes. The column temperature was maintained at 60°C with an injection volume of 2.5µl.

### D- and L-amino Acids Extraction

For D- and L-amino acid quantification, a calibration curve was prepared with LC-MS grade water as solvent, including the following concentrations: 60 nM, 100 nM, 250 nM, 400 nM, 600 nM, 800 nM, 1000 nM, 2000 nM, 3000 nM, 5000 nM, 8000 nM, 10000 nM. QCs were prepared for the lowest concentration (QC1, 60 nM), intermediary (QC2, 2000 nM), and highest concentration (QC3, 10000 nM). The diastereoisomers were selected based on the linearity of the calibration curve R2 >0.99. D- and L-amino acids with R2 below 0.99 were excluded. A total of 100 μl of each cecum content sample (20x and 100x diluted samples), QCs and calibration curve points were added to a 1 ml 96-well plate and mixed with 25 μl of an internal standard solution containing 100 μM of each stable-isotope labelled amino acid. Both double (solvent only, no internal standard) and single (solvent and internal standard mix) blanks were added to the plate. Then, samples were mixed with 750 μl of cold acetonitrile and centrifuged at 10 min, 13000 x *g* at 4°C to remove any protein precipitate. The supernatant was collected and dried overnight in a vacuum concentrator. The residue was dissolved in 50 μl of LC-MS grade water, followed by the addition of 35 μl of 0.15 M sodium tetraborate and 50 μl of a 2.5 mg/ml (S)-NIFE solution in acetonitrile. After 10 min incubation at room temperature, the reaction was terminated by adding 10 μl of 4 M HCl and 355 μl of LC-MS grade water.

The diastereoisomers were separated on a Waters ACQUITY UPLC BEH C18 2.1 × 150 mm, 1.7 μm column using an Acquity UPLC system coupled to an XEVO TQ-S tandem mass spectrometer (Waters Co, Milford MA, USA). The column was maintained at a constant temperature of 60 °C, with a mobile phase of 10 mM ammonium hydrogen carbonate (pH 9.5) (mobile phase A) and acetonitrile (mobile phase B). The desolvation gas flow was set at 900 L/hr, the cone gas flow at 150 L/hr, and a temperature of 650 °C. The gradient separation was completed in 23 minutes with the initial conditions of 96 % of A and flow rate of 0.6 ml/min for 5 min. Then, solvent B was increased to 10 % for 4 min, increasing gradually to 11.5 % during 2 min, 28 % for 5 min, reaching 35 % with 21 minutes of run time. Then, the solvent A increases to 75 % for 0.5 min and returns to the initial condition for 2 minutes. The column temperature was maintained at 60°C with an injection volume of 5µl.

### Peak Integration and Data Normalization of Bile Acids And D- and L-amino Acids

Peak integration of the bile acid and D- & L-amino acid raw data was performed using TargetLynx Application Manager in Waters MassLynx Software (version 4.2, Waters Laboratory Informatics). Retention time and concentration of the internal standards were used as references to the identification and quantification of the metabolites. Data with R2 above 0.99 was accepted for further analysis. Data (in nM) with valid values were normalized by the initial faecal weight (mg/ml). Samples with missing values were replaced by a 1/5 minimum value, and the data was log transformed.

### Statistical Analyses

Comparison of colonization burden between groups over multiple time points was accomplished using 2-way ANOVA testing. Comparison between groups for colonization burden, bacterial growth density, amino acid and bile acid content at discrete time points was performed with Mann-Whitney U testing with correction for multiple simultaneous comparisons using a False Discovery Rate (FDR) of 10% with the 2 stage step-up method of Benjamin, Krieger and Yekutieli. Comparison of colonization incidence was performed with Fisher’s Exact Test. For correlation between cecum bile acid content and Kp175 burden line of best fit and R^2^ were calculated with least squares regression with Kp175 burden as logarithmic variable and bile acid content as linear variable. Correlations were calculated with Spearman method. GraphPad Prism 10 software was used for all statistical comparisons.

### Study Approval

Mouse experiments were performed in strict accordance with recommendations from the National Institutes of Health Guide for the Care and Use of Laboratory Animals under a protocol approved and overseen by IACUC of the University of North Carolina at Chapel Hill (protocol #23-129).

### Data Availability

All data presented in this manuscript will be made available in the Supporting Data Values file upon publication.

## Author Contributions

TH: Designing research studies, conducting experiments, acquiring and analyzing data, writing manuscript. VS: Conducting experiments, acquiring and analyzing data, assistance with writing manuscript. JX: Conducting experiments, acquiring data. MFL: Conducting experiments, acquiring data. AB: Conducting experiments, acquiring data. KW: Assistance with conducting experiments. KW: Assistance with conducting experiments. JWA: Assistance with designing research studies, analyzing data, critical manuscript review. KM: Provision of reagents and materials, critical manuscript review. ON: Provision of reagents and materials, critical manuscript review. JRS: Assistance with designing research studies, critical manuscript review. DvD: Assistance with designing research studies, provision of reagents and materials, critical manuscript review. LAB: Scientific direction, designing research studies, provision of reagents and materials, critical manuscript review and revision.

## Acknowledgements

Research activities, animals, reagents and supplies and were funded through the National Institute of Allergy and Infectious Diseases (NIAID) of the National Institutes of Health (NIH); #T32-AI007151 (TH), #1R01-AI151214-01A1 (LAB), #1R01-AI143910-01 (DVD). JRS is supported by NIHR Southampton Biomedical Research Centre; OTN is supported by the Singapore Ministry of Health’s National Medical Research Council Clinician Scientist Award - Senior Investigator (MOH-001763). Access to the UNC National Gnotobiotic Rodent Resource Center was possible through National Institute of Diabetes and Digestive and Kidney Diseases of the NIH (#P30-DK034987, PI Robert S. Sandler). Bacterial isolates identified as “ARLG” were provided by the Antibacterial Resistance Leadership Group Laboratory Center, which is supported by the NIAID of the NIH under Award Number UM1AI104681. The content is solely the responsibility of the authors and does not necessarily represent the official views of the National Institutes of Health. Figures were generated using GraphPad Prism 10 and Microsoft Powerpoint software. The contents of this article are solely the responsibility of the authors and do not necessarily represent the official views of the NIH.

## Supplemental Material

**Figure S1:**
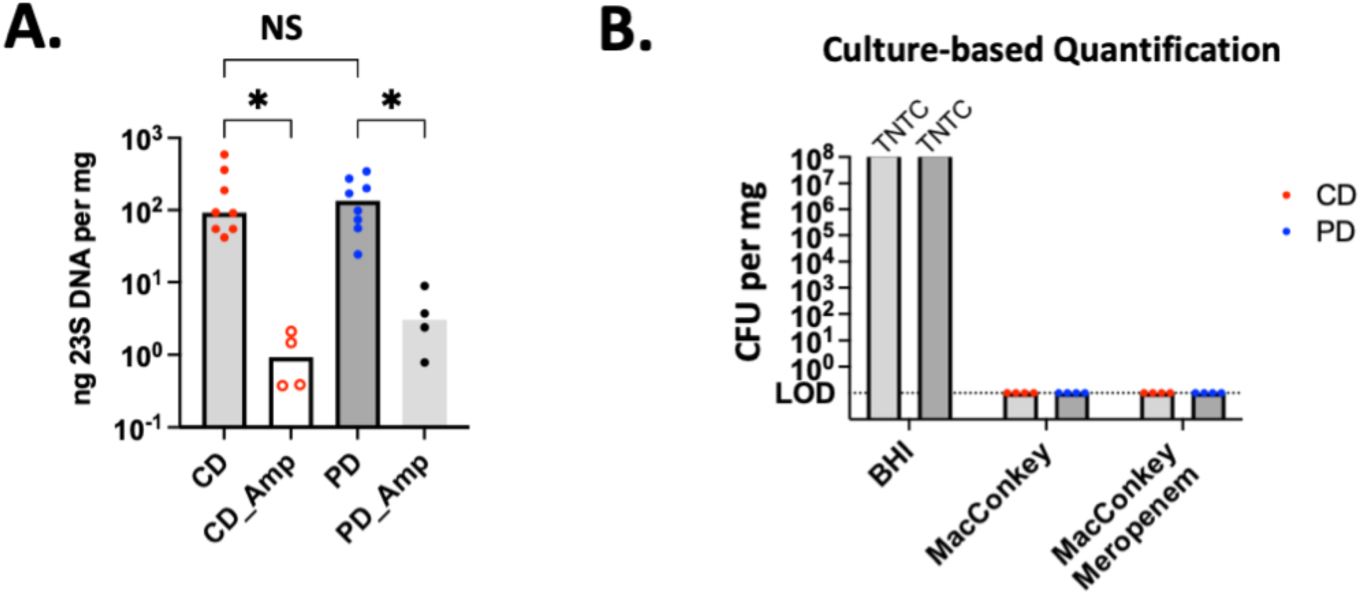
Quantification of Bacteria in Feces and Ceca of Mice fed CD or PD. (A.) Fecal specimens were collected from CD- and PD-fed mice, with or without ampicillin in their water for the previous 7 days. qPCR detection of 23S rRNA content was used for total bacterial burden quantification. Median concentration depicted, N=4-8 per group. Mann-Whitney U-test with 10% FDR performed, *= p<0.05, NS= No significance. (B.) Burden of aerobic bacteria expressed as log CFU per mg of cecal content growing on BHI, MacConkey, and MacConkey + 0.125 ug/ml meropenem agar in mice initiated on respective diets 7 days prior to sacrifice. Median concentration depicted. N = 4 per group. TNTC = too numerous to count.

**Figure S2:**
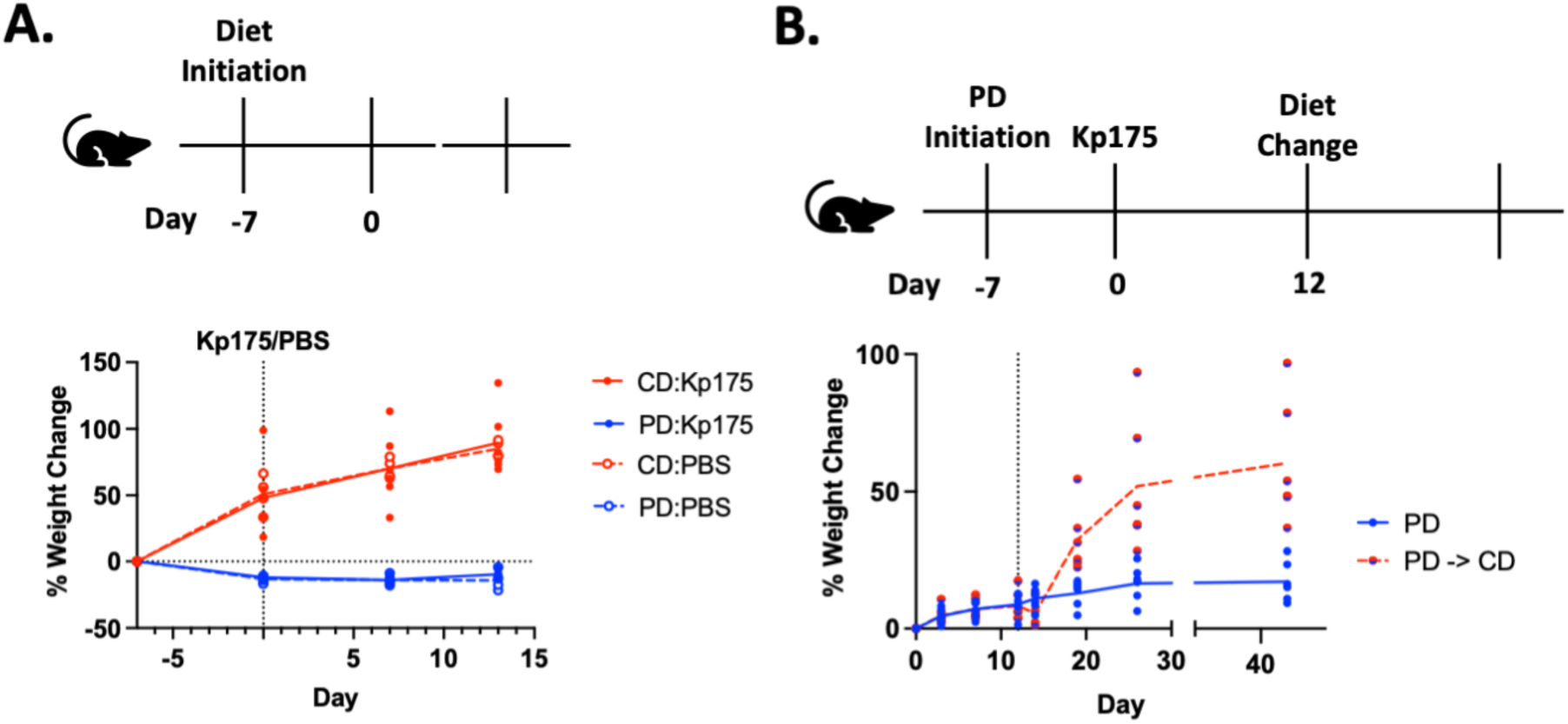
Dietary effect on mouse weight. (A.) Mice were initiated on CD or PD 7 days prior to oral gavage with 10^6^ CFU Kp175 or PBS, weight change expressed relative to day of diet initiation. N=4 per group. (B.) Mice were initiated on PD 7 days prior to oral gavage with 10^6^ CFU Kp175 or PBS, and in half of the mice diet switched to CD on day 12. Weight change expressed relative to day of inoculation. Mean weight change depicted, N=6 per group.

**Figure S3:**
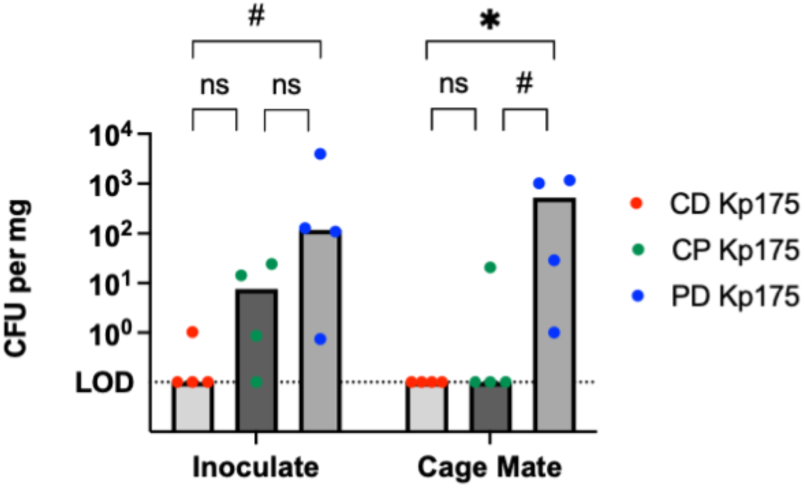
Peak Kp175 Burden and Transmission in Mice Fed CD, PD, or CP. 3-4 week old C57BL/6 mice placed on respective diets 7 days prior to inoculation with 10^6^ CFU Kp175 or PBS, with one Kp175 inoculated mouse and one PBS control cage-mate placed together in a single cage on day 2. Day 10 Kp175 burden expressed as log CFU per mg fecal specimen. Median burden depicted, N = 4 per group. Mann-Whitney U-test with 10% FDR performed, *= p<0.05, #= p<0.1, ns= no significance

**Figure S4:**
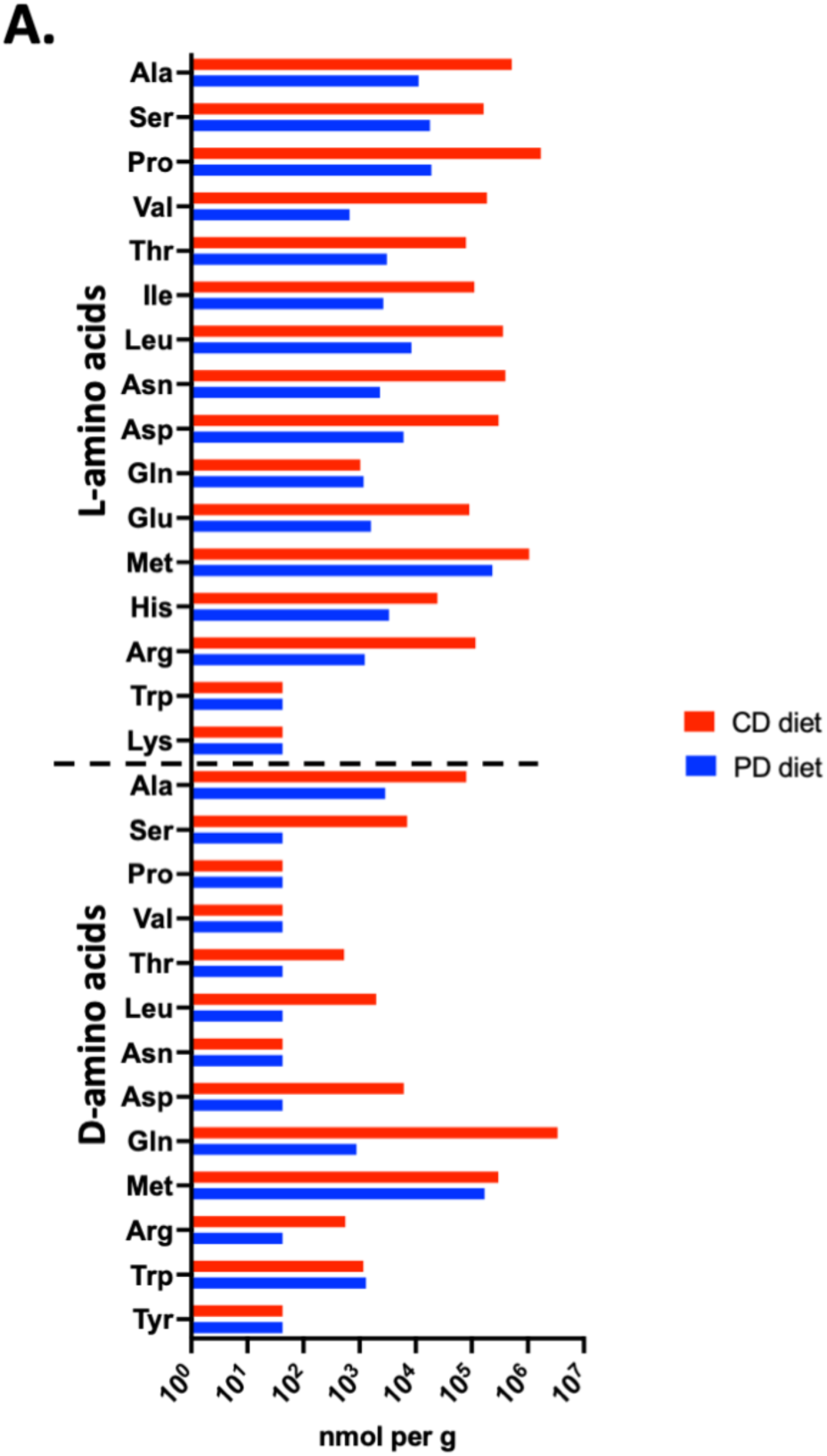
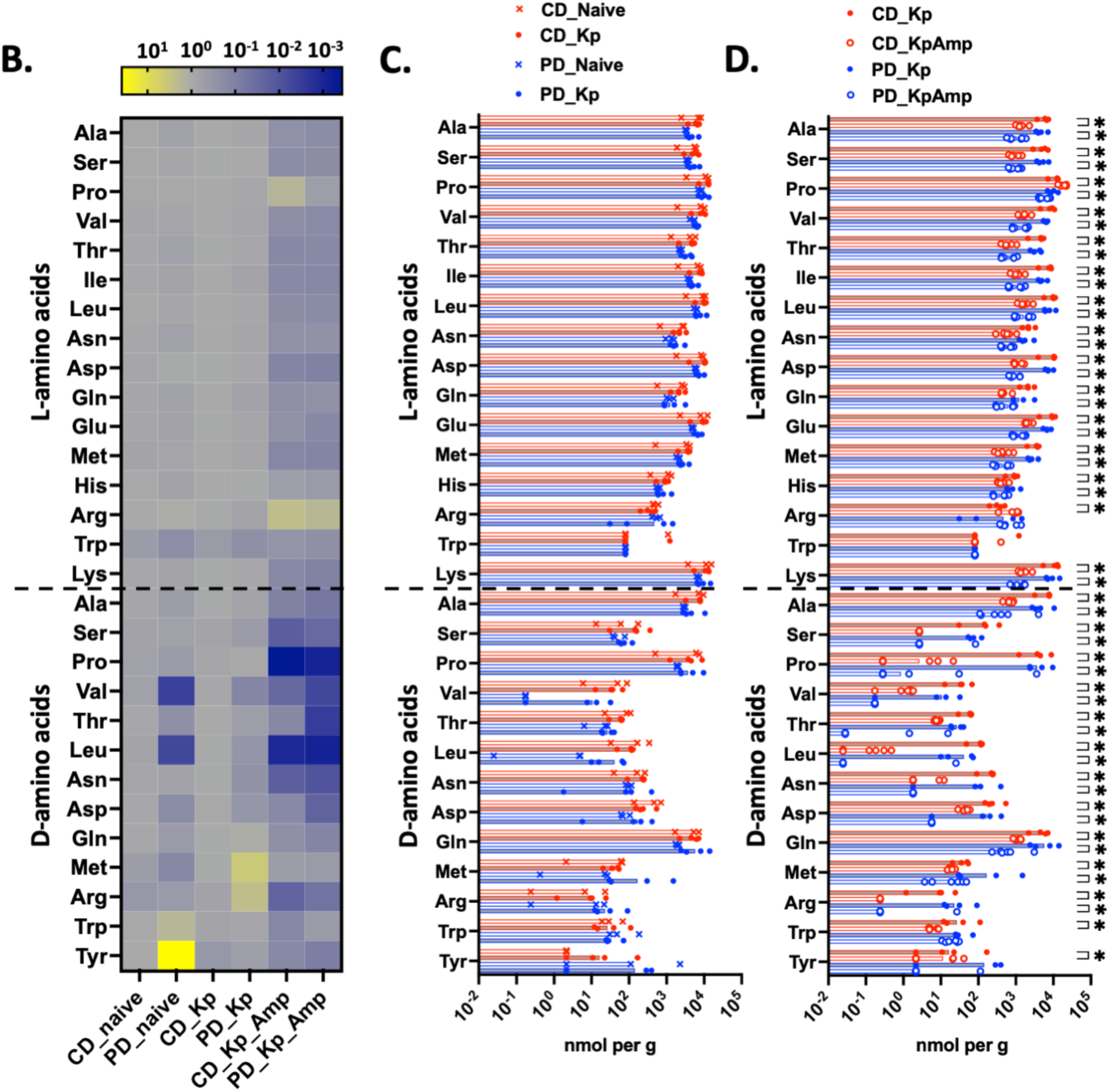
Amino Acid Content in Ceca of CD- and PD-fed Mice and in Mouse Food. (A.) Concentration of individual free L- and D-amino acids in CD or PD food. (B.) Heat map of individual free L- and D-amino acids in cecal content of mice placed on respective diets 7 days prior to inoculation with 10^6^ CFU Kp175 (or naïve without inoculation) and terminated on day 14 post-inoculation or treated with Ampicillin in the drinking water (1 mg/ml) at day 35 post-inoculation and terminated at day 42. Expressed as fold change relative to average concentration in cecal contents of colonized CD-fed mice without antibiotic treatment. Median concentration depicted, N=3-6 per group. (C.) Concentration of individual free L- and D-amino acids in cecal content of mice placed on respective diets 7 days prior to inoculation with 10^6^ CFU Kp175 or left without inoculation (naïve) and terminated on day 14 post-inoculation. Median concentration depicted, N=3-4 per group. Mann-Whitney U-test with 10% FDR performed. (D.) Concentration of individual free L- and D-amino acids in cecal content of mice placed on respective diets 7 days prior to inoculation with 10^6^ CFU Kp175 treated with Ampicillin in the drinking water (1 mg/ml) at day 35 post-inoculation and terminated at day 42. Median concentration depicted, N=4-6 per group, Mann-Whitney U-test with 10% FDR performed, *= p<0.05 by Mann-Whitney U-test.

**Figure S5:**
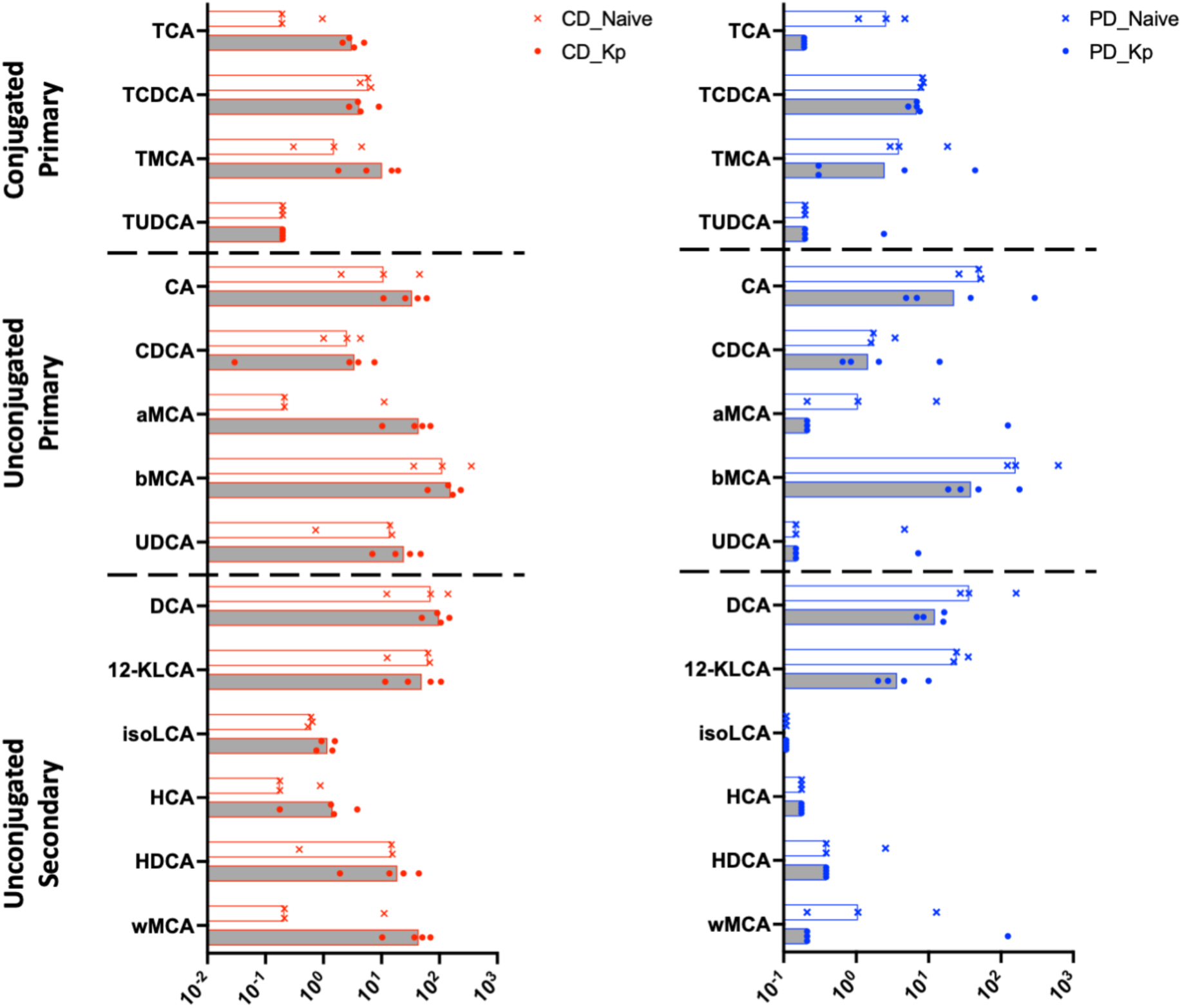
Bile Acid Content in Ceca of Mice With or Without Kp175 Colonization. Concentration of individual L- and D-amino acids in cecal content of mice placed on respective diets 7 days prior to oral gavage with 10^6^ CFU Kp175 or left without inoculation (naïve) and terminated on day 14 post-inoculation. Median concentration depicted, N=3-4 per group. Mann-Whitney U-test with 10% FDR performed. *TCA= Taurocholic Acid, TCDCA= Taurochenodeoxycholic Acid, TMCA= Tauromuricholic Acid, TUDCA= Tauroursodeoxycholic Acid, CA= Cholic Acid, CDCA= Chenodeoxycholic Acid, aMCA= Alpha Muricholic Acid, bMCA= Beta Muricholic Acid, UDCA= Ursodeoxycholic Acid, DCA= Deoxycholic Acid, 12-KLCA= 12-Ketolithocholic Acid, isoLCA= Isolithocholic Acid, HCA= Hyocholic Acid, HDCA= Hyodeoxycholic Acid, wMCA= Omega Muricholic Acid*.

**Table S1:**
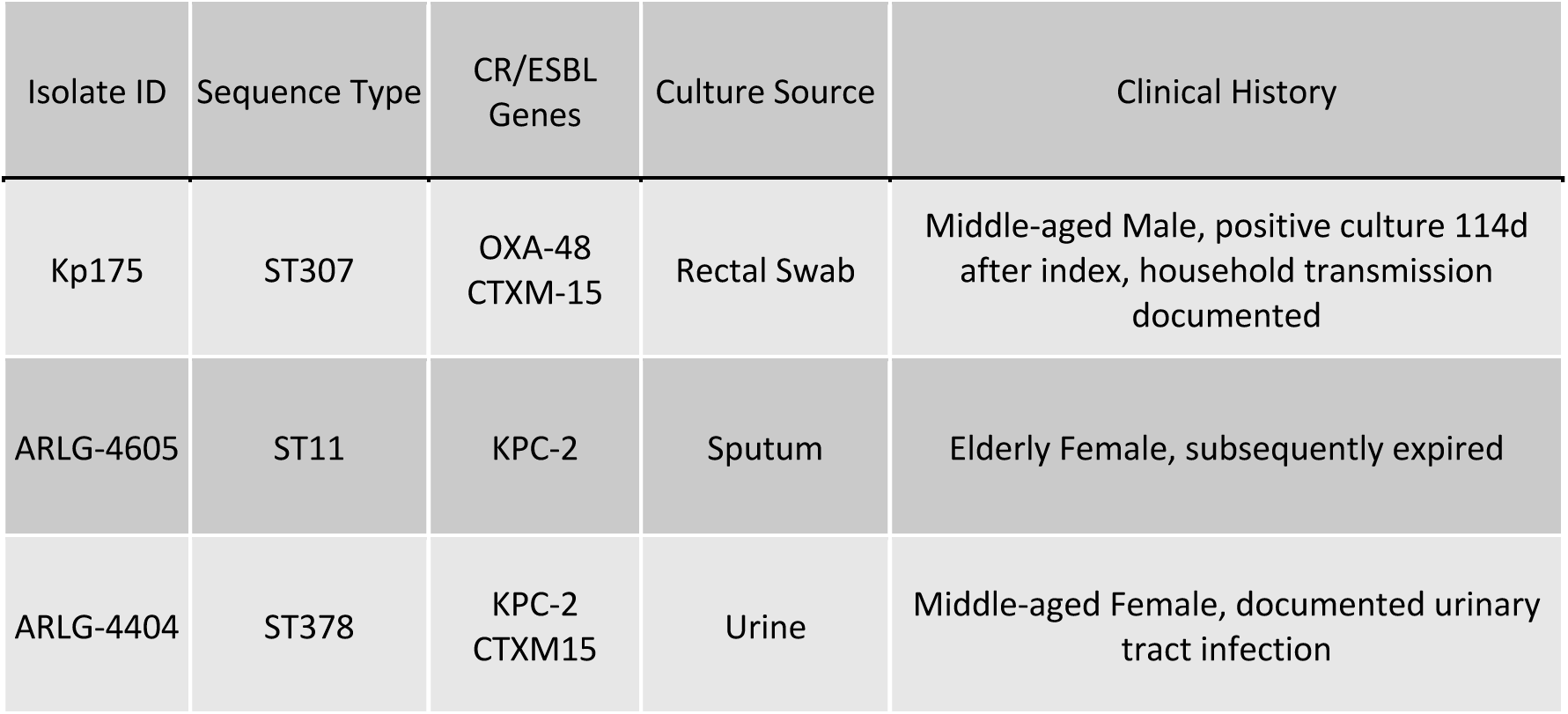
Carbapenem Resistant Klebsiella pneumoniae Isolates. Sequence type, antimicrobial resistance genes, and clinical features of isolates used in study.

| Isolate ID | Sequence Type | CR/ESBL Genes | Culture Source | Clinical History |
| --- | --- | --- | --- | --- |
| Kp175 | ST307 | OXA-48<br>CTXM-15 | Rectal Swab | Middle-aged Male, positive culture 114d after index, household transmission documented |
| ARLG-4605 | ST11 | KPC-2 | Sputum | Elderly Female, subsequently expired |
| ARLG-4404 | ST378 | KPC-2<br>CTXM15 | Urine | Middle-aged Female, documented urinary tract infection |

**Table S2:** Bile Acid Content in Ceca of Colonized Mice With or Without Ampicillin Treatment. Primary and secondary bile acid content and Kp175 burden compared from ceca of mice placed on respective diets 7 days prior to inoculation with 10^6^ CFU and terminated on day 14 post-inoculation or treated with Ampicillin in the drinking water (1 mg/ml) at day 35 post-inoculation and sacrificed at day 42. R^2^ calculated with least squares regression with X as logarithmic variable and Y as linear variable and r and p calculated using Spearman correlation method. *TCA= Taurocholic Acid, TCDCA= Taurochenodeoxycholic Acid, TMCA= Tauromuricholic Acid, TUDCA= Tauroursodeoxycholic Acid, CA= Cholic Acid, CDCA= Chenodeoxycholic Acid, aMCA= Alpha Muricholic Acid, bMCA= Beta Muricholic Acid, UDCA= Ursodeoxycholic Acid, DCA= Deoxycholic Acid, 12-KLCA= 12-Ketolithocholic Acid, isoLCA= Isolithocholic Acid, HCA= Hyocholic Acid, HDCA= Hyodeoxycholic Acid, wMCA= Omega Muricholic Acid*.

|  | R<sup>2</sup> | Pearson r | P value |
| --- | --- | --- | --- |
| TCA | 0.6301 | 0.726 | 0.0003 |
| TCDCA | 0.01925 | -0.4632 | 0.0397 |
| TMCA | 0.3116 | 0.6423 | 0.0023 |
| TUDCA | 0.4946 | 0.676 | 0.0011 |
| **Total Conjugated** | **0.6185** | 0.7534 | **0.0001** |
| CA | 0.0487 | -0.6762 | 0.0011 |
| CDCA | 0.1598 | -0.688 | 0.0008 |
| aMCA | 0.2434 | -0.6561 | 0.0017 |
| bMCA | 0.6264 | -0.7142 | 0.0004 |
| UDCA | 0.5751 | -0.6858 | 0.0008 |
| **Total Unconjugated** | **0.3518** | -0.7108 | **0.0004** |
| DCA | 0.7448 | -0.8605 | <0.0001 |
| 12-KLCA | 0.558 | -0.8707 | <0.0001 |
| isoLCA | 0.7437 | -0.6862 | 0.0008 |
| HCA | 0.4066 | -0.5418 | 0.0136 |
| HDCA | 0.4693 | -0.684 | 0.0009 |
| wMCA | 0.2434 | -0.6561 | 0.0017 |
| **Total Secondary** | **0.6606** | -0.8503 | **<0.0001** |

## References

1. Collaborators GBDAR. Global burden of bacterial antimicrobial resistance 1990-2021: a systematic analysis with forecasts to 2050. Lancet. 2024;404(10459):1199–226.

2. Antimicrobial Resistance Collaborators. Global burden of bacterial antimicrobial resistance in 2019: a systematic analysis. Lancet. 2022;399:629–55.

3. Shimasaki T, Seekatz A, Bassis C, Rhee Y, Yelin RD, Fogg L, et al. Increased Relative Abundance of Klebsiella pneumoniae Carbapenemase-producing Klebsiella pneumoniae Within the Gut Microbiota Is Associated With Risk of Bloodstream Infection in Long-term Acute Care Hospital Patients. Clin Infect Dis. 2019;68(12):2053–9.

4. Taur Y, Xavier JB, Lipuma L, Ubeda C, Goldberg J, Gobourne A, et al. Intestinal domination and the risk of bacteremia in patients undergoing allogeneic hematopoietic stem cell transplantation. Clin Infect Dis. 2012;55(7):905–14.

5. Gorrie CL, Mirceta M, Wick RR, Edwards DJ, Thomson NR, Strugnell RA, et al. Gastrointestinal Carriage Is a Major Reservoir of Klebsiella pneumoniae Infection in Intensive Care Patients. Clin Infect Dis. 2017;65(2):208–15.

6. Bar-Yoseph H, Hussein K, Braun E, and Paul M. Natural history and decolonization strategies for ESBL/carbapenem-resistant Enterobacteriaceae carriage: systematic review and meta-analysis. J Antimicrob Chemother. 2016;71(10):2729–39.

7. Marimuthu K, Mo Y, Ling ML, Hernandez-Koutoucheva A, Fenlon SN, Bertrand D, et al. Household transmission of carbapenemase-producing Enterobacteriaceae: a prospective cohort study. J Antimicrob Chemother. 2021;76(5):1299–302.

8. Ben-Ami R, Rodriguez-Bano J, Arslan H, Pitout JD, Quentin C, Calbo ES, et al. A multinational survey of risk factors for infection with extended-spectrum beta-lactamase-producing enterobacteriaceae in nonhospitalized patients. Clin Infect Dis. 2009;49(5):682–90.

9. Gasink LB, Edelstein PH, Lautenbach E, Synnestvedt M, and Fishman NO. Risk factors and clinical impact of Klebsiella pneumoniae carbapenemase-producing K. pneumoniae. Infect Control Hosp Epidemiol. 2009;30(12):1180–5.

10. Lim CJ, Cheng AC, Kennon J, Spelman D, Hale D, Melican G, et al. Prevalence of multidrug-resistant organisms and risk factors for carriage in long-term care facilities: a nested case-control study. J Antimicrob Chemother. 2014;69(7):1972–80.

11. Antimicrobial Resistance C. The burden of bacterial antimicrobial resistance in the WHO African region in 2019: a cross-country systematic analysis. Lancet Glob Health. 2024;12(2):e201–e16.

12. WHO. Malnutrition. https://www.who.int/news-room/fact-sheets/detail/malnutrition. Updated December 20, 2023 Accessed January 25, 2024.

13. Holowka T, van Duin D, and Bartelt LA. Impact of childhood malnutrition and intestinal microbiota on MDR infections. JAC Antimicrob Resist. 2023;5(2):dlad051.

14. Ndir A, Diop A, Faye PM, Cisse MF, Ndoye B, and Astagneau P. Epidemiology and Burden of Bloodstream Infections Caused by Extended-Spectrum Beta-Lactamase Producing Enterobacteriaceae in a Pediatric Hospital in Senegal. PLoS One. 2016;11(2):e0143729.

15. Berkley JA, Lowe BS, Mwangi I, Williams T, Bauni E, Mwarumba S, et al. Bacteremia among children admitted to a rural hospital in Kenya. N Engl J Med. 2005;352(1):39–47.

16. Andersen CT, Langendorf C, Garba S, Sayinzonga-Makombe N, Mambula C, Mouniaman I, et al. Risk of community- and hospital-acquired bacteremia and profile of antibiotic resistance in children hospitalized with severe acute malnutrition in Niger. Int J Infect Dis. 2022;119:163–71.

17. Woerther PL, Angebault C, Jacquier H, Hugede HC, Janssens AC, Sayadi S, et al. Massive increase, spread, and exchange of extended spectrum beta-lactamase-encoding genes among intestinal Enterobacteriaceae in hospitalized children with severe acute malnutrition in Niger. Clin Infect Dis. 2011;53(7):677–85.

18. Maataoui N, Langendorf C, Berthe F, Bayjanov JR, van Schaik W, Isanaka S, et al. Increased risk of acquisition and transmission of ESBL-producing Enterobacteriaceae in malnourished children exposed to amoxicillin. J Antimicrob Chemother. 2020;75(3):709–17.

19. . WHO guideline on the prevention and management of wasting and nutritional oedema (acute malnutrition) in infants and children under 5 years. Geneva; 2023.

20. Ling W, Peri AM, Furuya-Kanamori L, Harris PNA, and Paterson DL. Carriage Duration and Household Transmission of Enterobacterales Producing Extended-Spectrum Beta-Lactamase in the Community: A Systematic Review and Meta-Analysis. Microb Drug Resist. 2022;28(7):795–805.

21. Spragge F, Bakkeren E, Jahn MT, E BNA, Pearson CF, Wang X, et al. Microbiome diversity protects against pathogens by nutrient blocking. Science. 2023;382(6676):eadj3502.

22. Yip AYG, King OG, Omelchenko O, Kurkimat S, Horrocks V, Mostyn P, et al. Antibiotics promote intestinal growth of carbapenem-resistant Enterobacteriaceae by enriching nutrients and depleting microbial metabolites. Nat Commun. 2023;14(1):5094.

23. van der Waaij D, Berghuis-de Vries JM, and Lekkerkerk L-v. Colonization resistance of the digestive tract in conventional and antibiotic-treated mice. J Hyg (Lond). 1971;69(3):405–11.

24. Buffie CG, and Pamer EG. Microbiota-mediated colonization resistance against intestinal pathogens. Nat Rev Immunol. 2013;13(11):790–801.

25. Caballero-Flores G, Pickard JM, and Nunez G. Microbiota-mediated colonization resistance: mechanisms and regulation. Nat Rev Microbiol. 2023;21(6):347–60.

26. Joseph L, Merciecca T, Forestier C, Balestrino D, and Miquel S. From Klebsiella pneumoniae Colonization to Dissemination: An Overview of Studies Implementing Murine Models. Microorganisms. 2021;9(6).

27. Kizilates F, Yakupogullari Y, Berk H, Oztoprak N, and Otlu B. Risk factors for fecal carriage of extended-spectrum beta-lactamase-producing and carbapenem-resistant Escherichia coli and Klebsiella pneumoniae strains among patients at hospital admission. Am J Infect Control. 2021;49(3):333–9.

28. Young TM, Bray AS, Nagpal RK, Caudell DL, Yadav H, and Zafar MA. Animal Model To Study Klebsiella pneumoniae Gastrointestinal Colonization and Host-to-Host Transmission. Infect Immun. 2020;88(11).

29. Bartelt LA, Bolick DT, Mayneris-Perxachs J, Kolling GL, Medlock GL, Zaenker EI, et al. Cross-modulation of pathogen-specific pathways enhances malnutrition during enteric co-infection with Giardia lamblia and enteroaggregative Escherichia coli. PLoS Pathog. 2017;13(7):e1006471.

30. Brown EM, Wlodarska M, Willing BP, Vonaesch P, Han J, Reynolds LA, et al. Diet and specific microbial exposure trigger features of environmental enteropathy in a novel murine model. Nat Commun. 2015;6:7806.

31. Moore JH, Pinheiro CC, Zaenker EI, Bolick DT, Kolling GL, van Opstal E, et al. Defined Nutrient Diets Alter Susceptibility to Clostridium difficile Associated Disease in a Murine Model. PLoS One. 2015;10(7):e0131829.

32. Bartelt LA, Bolick DT, Kolling GL, Roche JK, Zaenker EI, Lara AM, et al. Cryptosporidium Priming Is More Effective than Vaccine for Protection against Cryptosporidiosis in a Murine Protein Malnutrition Model. PLoS Negl Trop Dis. 2016;10(7):e0004820.

33. Mayneris-Perxachs J, Bolick DT, Leng J, Medlock GL, Kolling GL, Papin JA, et al. Protein- and zinc-deficient diets modulate the murine microbiome and metabolic phenotype. Am J Clin Nutr. 2016;104(5):1253–62.

34. Bhatt AP, Arnold JW, Awoniyi M, Sun S, Santiago VF, Quintela PH, et al. Giardia Antagonizes Beneficial Functions of Indigenous and Therapeutic Intestinal Bacteria during Malnutrition. bioRxiv. 2024.

35. Giallourou N, Arnold J, McQuade ETR, Awoniyi M, Becket RVT, Walsh K, et al. Giardia hinders growth by disrupting nutrient metabolism independent of inflammatory enteropathy. Nat Commun. 2023;14(1):2840.

36. Bartelt LA, Roche J, Kolling G, Bolick D, Noronha F, Naylor C, et al. Persistent G. lamblia impairs growth in a murine malnutrition model. J Clin Invest. 2013;123(6):2672–84.

37. Bolick DT, Roche JK, Hontecillas R, Bassaganya-Riera J, Nataro JP, and Guerrant RL. Enteroaggregative Escherichia coli strain in a novel weaned mouse model: exacerbation by malnutrition, biofilm as a virulence factor and treatment by nitazoxanide. J Med Microbiol. 2013;62(Pt 6):896–905.

38. Bhatt AP, Arnold JW, Awoniyi M, Sun S, Feijoli Santiago V, Coskuner D, et al. Giardia antagonizes beneficial functions of indigenous and therapeutic intestinal bacteria during protein deficiency. Gut Microbes. 2024;16(1):2421623.

39. Larabi AB, Masson HLP, and Baumler AJ. Bile acids as modulators of gut microbiota composition and function. Gut Microbes. 2023;15(1):2172671.

40. Pi Y, Mu C, Gao K, Liu Z, Peng Y, and Zhu W. Increasing the Hindgut Carbohydrate/Protein Ratio by Cecal Infusion of Corn Starch or Casein Hydrolysate Drives Gut Microbiota-Related Bile Acid Metabolism To Stimulate Colonic Barrier Function. mSystems. 2020;5(3).

41. Ridlon JM, and Gaskins HR. Another renaissance for bile acid gastrointestinal microbiology. Nat Rev Gastroenterol Hepatol. 2024;21(5):348–64.

42. Kwong LH, Ercumen A, Pickering AJ, Arsenault JE, Islam M, Parvez SM, et al. Ingestion of Fecal Bacteria along Multiple Pathways by Young Children in Rural Bangladesh Participating in a Cluster-Randomized Trial of Water, Sanitation, and Hygiene Interventions (WASH Benefits). Environ Sci Technol. 2020;54(21):13828–38.

43. Sisay A, Asmare Z, Kumie G, Gashaw Y, Getachew E, Ashagre A, et al. Prevalence of carbapenem-resistant gram-negative bacteria among neonates suspected for sepsis in Africa: a systematic review and meta-analysis. BMC Infect Dis. 2024;24(1):838.

44. Crichton M, Craven D, Mackay H, Marx W, de van der Schueren M, and Marshall S. A systematic review, meta-analysis and meta-regression of the prevalence of protein-energy malnutrition: associations with geographical region and sex. Age Ageing. 2019;48(1):38–48.

45. Han JH, Lapp Z, Bushman F, Lautenbach E, Goldstein EJC, Mattei L, et al. Whole-Genome Sequencing To Identify Drivers of Carbapenem-Resistant Klebsiella pneumoniae Transmission within and between Regional Long-Term Acute-Care Hospitals. Antimicrob Agents Chemother. 2019;63(11).

46. Chan YQ, Chen K, Chua GT, Wu P, Tung KTS, Tsang HW, et al. Risk factors for carriage of antimicrobial-resistant bacteria in community dwelling-children in the Asia-Pacific region: a systematic review and meta-analysis. JAC Antimicrob Resist. 2022;4(2):dlac036.

47. Hecht AL, Harling LC, Friedman ES, Tanes C, Lee J, Firrman J, et al. Dietary carbohydrates regulate intestinal colonization and dissemination of Klebsiella pneumoniae. J Clin Invest. 2024;134(9).

48. Desai MS, Seekatz AM, Koropatkin NM, Kamada N, Hickey CA, Wolter M, et al. A Dietary Fiber-Deprived Gut Microbiota Degrades the Colonic Mucus Barrier and Enhances Pathogen Susceptibility. Cell. 2016;167(5):1339–53 e21.

49. Djukovic A, Garzon MJ, Canlet C, Cabral V, Lalaoui R, Garcia-Garcera M, et al. Lactobacillus supports Clostridiales to restrict gut colonization by multidrug-resistant Enterobacteriaceae. Nat Commun. 2022;13(1):5617.

50. Shen M, Zhao H, Han M, Su L, Cui X, Li D, et al. Alcohol-induced gut microbiome dysbiosis enhances the colonization of Klebsiella pneumoniae on the mouse intestinal tract. mSystems. 2024;9(3):e0005224.

51. Zhang L, Voskuijl W, Mouzaki M, Groen AK, Alexander J, Bourdon C, et al. Impaired Bile Acid Homeostasis in Children with Severe Acute Malnutrition. PLoS One. 2016;11(5):e0155143.

52. Zhao X, Setchell KDR, Huang R, Mallawaarachchi I, Ehsan L, Dobrzykowski Iii E, et al. Bile Acid Profiling Reveals Distinct Signatures in Undernourished Children with Environmental Enteric Dysfunction. J Nutr. 2021;151(12):3689–700.

53. Ibrahim MK, Zambruni M, Melby CL, and Melby PC. Impact of Childhood Malnutrition on Host Defense and Infection. Clin Microbiol Rev. 2017;30(4):919–71.

54. Kotloff KL, Nataro JP, Blackwelder WC, Nasrin D, Farag TH, Panchalingam S, et al. Burden and aetiology of diarrhoeal disease in infants and young children in developing countries (the Global Enteric Multicenter Study, GEMS): a prospective, case-control study. Lancet. 2013;382(9888):209-22.

55. Page AL, de Rekeneire N, Sayadi S, Aberrane S, Janssens AC, Rieux C, et al. Infections in children admitted with complicated severe acute malnutrition in Niger. PLoS One. 2013;8(7):e68699.

56. Holowka TA, A.; Khuu, T.; Bimagambetov, A.; Dial, C. N.; Kondwani, A.; Tepeka, A.; Alby, K.; Juliano, J. J.; Garcia-Pratts, A. J.; Mvalo, T.; Vonasek, B. J.; Ciccone, E. J.; Bartelt, L. A. Pervasive Intestinal Carriage with Multiple Species of Extended Spectrum Cephalosporin-Resistant Enterobacterales in Children Admitted for Severe Acute Malnutrition at a Tertiary Hospital in Malawi. medRxiv.

57. Bray AS, and Zafar MA. Deciphering the gastrointestinal carriage of Klebsiella pneumoniae. Infect Immun. 2024;92(9):e0048223.

58. Sun Y, Patel A, SantaLucia J, Roberts E, Zhao L, Kaye K, et al. Measurement of Klebsiella Intestinal Colonization Density To Assess Infection Risk. mSphere. 2021;6(3):e0050021.

